# Starvation sensing by mycobacterial RelA/SpoT homologue through constitutive surveillance of translation

**DOI:** 10.1101/2022.12.29.522164

**Authors:** Yunlong Li, Soneya Majumdar, Ryan Treen, Manjuli R. Sharma, Jamie Corro, Howard B. Gamper, Swati R. Manjari, Jerome Prusa, Nilesh K. Banavali, Christina L. Stallings, Ya-Ming Hou, Rajendra K. Agrawal, Anil K. Ojha

## Abstract

The stringent response, which leads to persistence of nutrient-starved mycobacteria, is induced by activation of the RelA/SpoT homologue (Rsh) upon entry of a deacylated-tRNA in a translating ribosome. However, the mechanism by which Rsh identifies such ribosomes *in vivo* remains unclear. Here, we show that conditions inducing ribosome hibernation result in loss of intracellular Rsh in a Clp protease-dependent manner. This loss is also observed in non-starved cells using mutations in Rsh that block its interaction with the ribosome, indicating that Rsh association with the ribosome is important for Rsh stability. The cryo-EM structure of the Rsh-bound 70S ribosome in a translation initiation complex reveals unknown interactions between the ACT domain of Rsh and components of the ribosomal L7/L12-stalk base, suggesting that the aminoacylation status of A-site tRNA is surveyed during the first cycle of elongation. Altogether, we propose a surveillance model of Rsh activation that originates from its constitutive interaction with the ribosomes entering the translation cycle.

**Significance:** Bacteria persist under nutrient starvation by activating RelA/SpoT homologue (Rsh), which synthesizes a growth regulating alarmone, ppGpp. Rsh is activated specifically upon recognizing a translation elongation complex with deacylated tRNA at the A-site. It is however unclear how Rsh identifies such a complex in vivo. We show here that conditions inducing ribosome hibernation in mycobacteria cause loss of intracellular Rsh, implying that association with translating ribosomes is necessary for intracellular stability of Rsh. Using structural analysis of Rsh-bound 70S translation initiation complex, we propose here that mycobacterial Rsh identifies a Rsh-activating ribosomal complex by constitutively surveying the ribosome entering the translation cycle at the early elongation stage.

## Introduction

The stringent response in bacteria is a programmed adaptation to nutrient starvation facilitated by the secondary messengers, guanosine tetra- or penta-phosphates [collectively called (p)ppGpp] (1–4). This response promotes long-term survival and persistence of mycobacteria under various stresses including antibiotics (5–10). The primary function of (p)ppGpp is to reprogram the global gene expression in cells either directly by binding to RNA polymerase (3, 11) or by binding to other cellular targets (12–14).

The level of (p)ppGpp is primarily regulated by a GDP/GTP pyrophosphokinase and a GDP/GTP pyrophosphohydrolase, encoded either as two enzymes RelA and SpoT in β- and γ-proteobacteria (3), or as a single protein called Rel in other species (15). Further insights into the domain architectures of RelA/Rel and the mechanism of its activation have been obtained from the structures of either the free enzyme or in complex with the ribosome, mRNA and A- and P-site tRNAs (16–20). RelA/Rel has five distinct domains (Fig. 1A): i) N-terminal domain possessing the (p)ppGpp synthase and hydrolase activities, ii) TGS (ThrRS, GTPase, SpoT) domain that interacts with 3’ CCA tail of deacylated-tRNA, iii) alpha helical domain (AHD), iv) zinc-binding domain (ZBD) that contacts the A-site finger (ASF) on the 23S rRNA and S19 at the inter-subunit interface, and v) the ACT domain (Aspartate Kinase, Chorismate and TyrA) occupying a cavity in the large 50S ribosomal subunit (or LSU) (17–19). The interaction between the TGS domain and 3’ CCA tail of the deacylated-tRNA stabilizes the tRNA in a distorted state at the A-site, called the A/R state, while pinning the domain against the small 30S ribosomal subunit (or SSU) and extending the flexible catalytic N-terminal domain of RelA in an open and active conformation (17–19).

**Figure 1:**
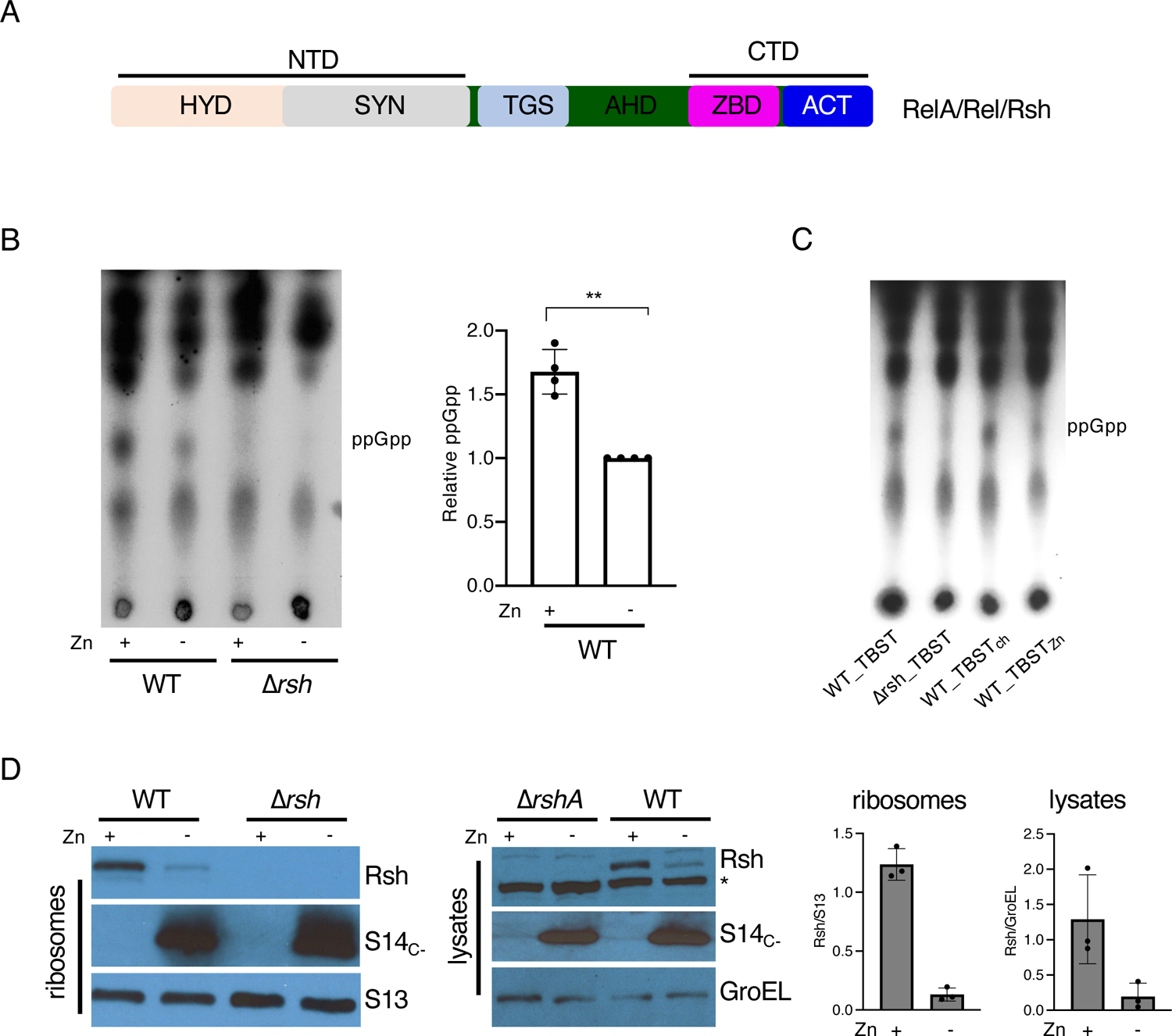
Attenuated stringent response and loss of Rsh in zinc-starved mycobacteria. **A.** A conserved domain architecture in mycobacterial RelA/SpoT homologue (Rsh) compared to RelA nd Rel in other species. The N-terminal domain (NTD) containing the hydrolase (HYD) and synthase (SYN) is distinguished from the C-terminal domain (CTD) by TGS (Thr-RS, GTPase and SpoT) and alpha-helix domain (AHD). The CTD is comprised of ZBD (zinc binding domain) and ACT (aspartokinase, chorismate mutase and TyrA) subdomains. **B.** Radio-TLC showing ppGpp levels in cultures of wild-type (WT) *M. smegmatis* isolated from high-zinc (1mM ZnSO_4_) and low-zinc (1 µM TPEN) Sauton’s medium after 96 hours of growth. ^32^P-labeled sodium phosphate (100 µCi/mL) was added to the cultures six hours prior to starvation in TBST for three hours. After starvation nucleotides were extracted in formic acid and amounts equivalent to 50000 cpm were analyzed by TLC. Purified cold ppGpp was loaded as positional marker. The plot shows ppGpp levels in high-zinc WT samples relative to their low-zinc counterparts from four biologically independent experiments. **C.** Radio-TLC showing ppGpp levels in WT cells cultured for 96-hours in high-zinc Sauton’s medium, then labelled for six hours, and resuspended in either TBST, or Chelex 100-treated TBST (TBST_ch_), or zinc-supplemented TBST_ch_ (TBST_Zn_). A Δ*rsh* mutant was used as control in panels B and C. **D**. Immunoblot analysis of Rsh levels in 70S ribosomes purified from 96-hour-old high-and low-zinc cultures of *M. smegmatis* strains described in panel B. S14c-was used as a marker for the remodeled C-ribosome, while S13 was used as the loading control for ribosomes. In the set of immunoblots on the right, cell lysates from the samples used for ribosome analysis were probed. The asterisk indicates a non-specific unknown protein cross-reactive to Rsh antibody. The plots to the extreme right show the normalized average difference in Rsh between high- and low-zinc cultures from three independent experiments.

The mechanism by which RelA finds a stalled ribosome complex in a nutrient starved cell is unclear. In the currently adopted ‘hopping model’, diffusible RelA samples translation elongation complexes and binds to a stalled ribosome with deacylated A-site tRNA in the A/R state, and upon catalyzing (p)ppGpp synthesis the enzyme ‘hops’ to another stalled complex (21–23). The hopping model further hypothesizes that an inactive RelA in non-starved cells acquires a ribosome-free, closed conformation, in which the catalytic activity is autoinhibited by the CTD, either through intermolecular (24–26) or intramolecular interactions (27–32). Winther et al. further proposed that free cytosolic RelA in *E. coli* binds to deacylated-tRNA before binding to the translating ribosome (22), although this model is challenged by the evidence of RelA bound to the ribosome without any tRNA at the A-site (18). Moreover, single-molecule tracking of RelA-fluorescent protein suggests constitutive association between RelA and ribosome in non-starved *E. coli* cells (33). Recognizing these discrepancies, Takada et al. offered a refined hopping model in *B. subtilis* in which activation of the stringent response appears to be initiated by a weak interaction between Rel and the ribosome before the complex is stabilized by the entry of deacylated-tRNA at the A-site, although they found RelA-ribosome interaction in *E. coli* to be too weak to universally validate their model (23). These studies while revealing subtle differences in the properties of RelA homologues across bacterial species, also underscore the limitations of biochemical assays in modeling a highly dynamic *in vivo* interaction between the stringent factor and the translation machinery.

Mycobacteria encode a single dual-function enzyme, RelA/SpoT homologue (henceforth called Rsh), with both kinase and hydrolase activities located in distinct domains of the enzyme (25, 26, 34–36). The optimal kinase activity of mycobacterial Rsh is observed in a Rsh Activating Complex (RAC) comprising mRNA, deacylated-tRNA and the ribosome (37). The ribosome in *Mycobacterium smegmatis* and *Mycobacterium tuberculosis* (Mtb) is remodeled under relatively moderate zinc starvation, and subsequently targeted for hibernation under severe zinc starvation (38, 39). Ribosome remodeling involves replacement of ribosomal proteins containing the CXXC motif (called C+ r-proteins) with their paralogs without the motif (called C-r-proteins) (38, 40). The C-ribosome is subsequently targeted for hibernation by mycobacterial protein Y (Mpy), with assistance from a zinc-sensing protein called Mpy recruitment factor (Mrf) (38, 39). A zinc-bound form of Mrf is sensitive to degradation by Clp protease, such that Mrf accumulation increases with decreasing concentration of zinc (39).

Investigating the interplay between the stringent response and ribosome hibernation in mycobacteria, we observed that interaction with the ribosome is necessary for the stability of intracellular Rsh in non-starved cells. The structure of Rsh-bound 70S ribosome in a translation initiation complex allows us to propose a new mechanism of starvation sensing by mycobacterial Rsh, in which its activation arises from its unique interaction with the ribosome entering the translation cycle.

## Results

### Mrf attenuates the stringent response in zinc-starved mycobacteria

We tested how a zinc-limiting growth condition that induces ribosome hibernation in *M. smegmatis* would impact ppGpp synthesis. After 96-hours of growth in high- or low-zinc Sauton’s medium, wild-type *M. smegmatis* cells were exposed for 3 hours to macronutrient starvation in tris-buffered saline with Tween-80 (TBST) to induce the stringent response to a measurable level (6). Rsh-dependent accumulation of ppGpp upon TBST starvation of the high-zinc culture was significantly greater than the low-zinc culture (Fig. 1B). To distinguish the contribution of macronutrient depletion from zinc limitation in TBST, we examined the stringent response in cells from high-zinc cultures after exposure to metal-chelated or zinc-supplemented TBST. Metal chelation did not impact the ppGpp synthesis, indicating that macronutrient starvation in TBST was the primary inducer of the stringent response (Fig. 1C).

Interaction with ribosomes is critical for the stringent response (21), we therefore hypothesized that the attenuated stringent response in low-zinc cultures was likely due to zinc-responsive changes in the ribosome structure and/or function. We therefore analyzed the Rsh occupancy on the ribosome in high- and low-zinc cultures. Rsh occupancy on the ribosomes purified from low-zinc cells was significantly lower than that from high-zinc cells (Fig. 1D). Unexpectedly, the difference in Rsh abundance on the ribosome correlated tightly with the levels in the total cellular lysate (Fig. 1D). Moreover, lack of any other detectable smaller Rsh isoforms (Fig S1) suggests total loss of intracellular Rsh. Together, figures 1B-D show that physiological conditions developed during zinc starvation decreases the abundance of Rsh and thus its interaction with the ribosome, thereby diminishing the stringent response.

We next investigated the effect of two zinc-responsive physiological changes in the ribosome, remodeling and hibernation, in altered Rsh stability and Rsh-ribosome interaction. The Zur-mediated C+ to C-remodeling of the ribosome induced by zinc starvation did not impact the Rsh-ribosome interaction: Rsh occupancy on the ribosome from a high-zinc culture of wildtype was comparable to Δ*zur* and a strain constitutively synthesizing C-ribosomes (Fig. S2). To determine the effect of zinc-responsive ribosome hibernation on Rsh-ribosome interaction, we analyzed the impact of Mpy recruitment factor (Mrf) on Rsh stability, activity, and its occupancy on the ribosome. Mrf binds to SSU and facilitates inhibition of translation and growth retardation upstream to Mpy recruitment to the ribosome (39). In zinc-starved *M. smegmatis,* deletion of *mrf* restored the levels of intracellular and ribosome-bound Rsh similar to those in high-zinc wildtype cells, and the mutant phenotype was complemented by plasmid-borne *mrf* (Fig. 2A-B). The increase in Rsh abundance in zinc-starved Δ*mrf* also correlated with the increase in ppGpp levels upon exposure to TBST (Fig. 2C). Together, these data suggest that: i) the limiting intracellular free zinc concentration has little direct impact on the stringent response, and ii) accumulation of Mrf induced by zinc depletion and likely downregulation of translational activity decrease total Rsh abundance in the cell. Deletion of *rsh* did not impact Mpy-ribosome interaction (Fig. S3), distinguishing mycobacteria from other species in which ppGpp transcriptionally induces factors necessary for ribosome hibernation (41–43).

**Figure 2:**
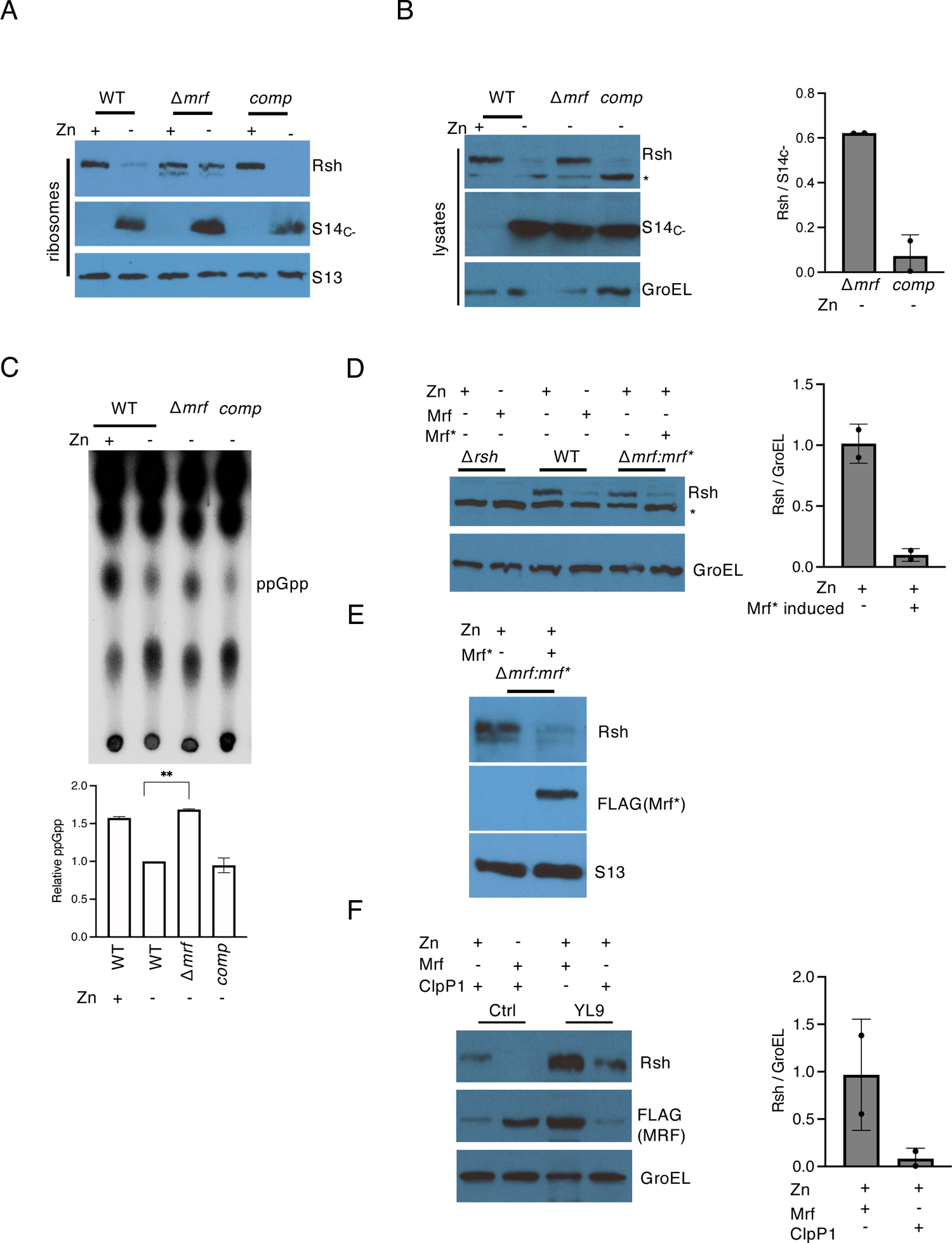
Mrf accumulation causes loss of Rsh. **A-B.** Analysis of Rsh in 70S ribosomes (A) and cell lysates (B) from high- or low-zinc cultures of Δ*mrf* and complemented (Δ*mrf*comp) strains of *M. smegmatis.* High and low-zinc cultures of WT were controls. S14_c-_, S13 and GroEL were probed as loading controls for ribosomes or cell lysates as indicated. The asterisk in panel B indicates a non-specific protein cross-reactive to Rsh antibody. **C.** Radio-TLC showing ppGpp levels in 96-hour old high- or low-zinc cultures of WT and low-zinc cultures of Δ*mrf and* Δ*mrfcomp* of *M. smegmatis* strains after 3-hours of starvation in TBST. Purified cold ppGpp was loaded as positional marker. The plot shows relative ppGpp levels in each sample compared to the level in low-zinc culture of WT from two biologically independent experiments. **D.** Ectopic Mrf expression causes loss of Rsh. Levels of Rsh in the cell lysates of a *M. smegmatis* strain (pYL222) expressing FLAG-tagged Mrf* under the acetamide-inducible promoter. Cells were cultured in high-zinc Sauton’s medium (1mM ZnSO4) containing 0.2% succinate as the carbon source until OD 0.6, after which Mrf* expression was induced with 0.2% acetamide, and cells were grown for 96 hours from inoculation. Uninduced cells were used as the reference. As controls, 96-hour old cells from the wildtype (WT) and Δ*rsh* strains cultured in either high- or low-zinc (1µM TPEN) Sauton’s medium with succinate as the carbon source. The plot shows average normalized density of Rsh from two independent experiments. **E.** Analysis of Rsh in ribosomes purified from 96-hour-old induced or uninduced high-zinc cultures of pYL222 strain. **F.** Simultaneous accumulation of Rsh and Mrf in YL9 cells (Li et al. 2020) upon conditional depletion of ClpP1 protease. Cells were grown in high-zinc Sauton’s medium until OD of 0.6, after which anhydrotetracycline (ATc) was added to induce CRISPRi system to deplete ClpP1. Cell lysates were analyzed after 16 hours of ClpP1 depletion. A parallel uninduced cells were used as control. YL9 cells grown for 96 hours in high- or low-zinc Sauton’s medium without ATc were used as controll. The plot shows average normalized density of Rsh from two independent experiments.

### Ectopic expression of Mrf can reduce intracellular abundance of Rsh

To further establish the relationship between Rsh abundance and Mrf accumulation, we examined the impact of an ectopic expression of a Mrf variant (Mrf*) on Rsh levels in zinc-rich cells. Mrf* has reduced zinc affinity and is recalcitrant to regulation by the Clp protease system (39). Consequently, constitutive expression of Mrf* leads to its substantial accumulation in high-zinc mycobacteria (39), presumably in a conformation that is capable of binding to the ribosome and inhibiting both translation and cellular growth. The abundance of Rsh and its association with the ribosome was determined upon acetamide-inducible expression of Mrf* in high-zinc cultures of *M. smegmatis*. Figure 2D shows that the expression of Mrf* in high-zinc cells led to concomitant loss of Rsh, both in the lysates (Fig. 2D) and on the ribosomes (Fig. 2E).

Because SpoT in *Salmonella enterica* was identified as a potential substrate of the Clp protease system (44), we tested the possibility whether mycobacterial ClpP1P2 protease could be involved in regulating the abundance of free Rsh in cells. We used a *M. smegmatis* strain, which constitutively transcribed FLAG-tagged Mrf and harbored an anhydrotetracycline-inducible CRISPR-Ca9 system for targeting ClpP1 (39), to test whether Rsh loss during Mrf stabilization is Clp-dependent. Depletion of ClpP1 in high-zinc cells resulted in concomitant accumulation of both Mrf and Rsh (Fig. 2F). Thus, loss of Rsh in cells during Mrf stabilization is likely achieved through a Clp-dependent mechanism, although whether Rsh is a direct substrate of Clp remains unknown.

### Interaction with the ribosome is necessary for intracellular stability of Rsh

Above, we have established loss of Rsh in cells with progressively increasing amounts of Mrf prior to ribosome hibernation, suggesting that availability of translating ribosome might be important for Rsh stability. Moreover, similar levels of Rsh-bound ribosomes in high-zinc cultures before and after nutrient starvation (Fig. S4), suggesting that there is a constant level of Rsh-ribosome interaction regardless of the changes in the relative levels of (p)ppGpp. We therefore tested whether interaction with the ribosome impacts the intracellular stability of Rsh. *E. coli* RelA interacts with the ribosome primarily through its CTD containing a conserved CCHC type zinc-finger in the ZBD, which makes extensive contacts with the inter-subunit region of the 70S ribosome, and ASF of the LSU (17, 45). Although the requirement of zinc in RelA-ribosome interaction remains unclear, the zinc coordinating residues C612, C613, H634 and C638 (C666, C667, H688 and C692 in *M. smegmatis Rsh*) of ZBD are conserved in mycobacteria and likely acquire the same three-dimensional conformation as in *E. coli* RelA (Fig. 3A). Furthermore, mutation of C633 in *M. tuberculosis* Rsh (C692 in *M. smegmatis*) impairs the enzyme *in vitro* (25), supporting an important role of this residue in the Rsh activity. We therefore mutated the key ZBD residues and studied the impact on the intracellular abundance of Rsh. The mutation scheme closely mirrored the previous study (24), which also showed that mutation in the neighboring D637 (D691 in *M. smegmatis*) had substantially less impact on protein function compared to C638 (C692 in *M. smegmatis*). Substitution of C666 and C692 residues resulted in substantial loss of Rsh in both the cell lysate and on the ribosome in actively growing *M. smegmatis* cells in zinc-rich conditions (Fig. 3B). The presence of wildtype Rsh on the ribosome was unambiguous (Fig. 3B), supporting the notion that Rsh-ribosome interaction in non-starved cells is constitutive. Substitution in the adjacent D691 neither impacted the Rsh-ribosome interaction nor the intracellular abundance of the protein (Fig. 3B). Recombinant expression of these Rsh variants in *E. coli* did not significantly impact the protein yield (Fig. S5), indicating that these mutations do not affect the intrinsic stability of the protein *in vivo*, and their properties are likely determined by the intracellular environment of mycobacteria. To further test if the mutations specifically disrupted Rsh-ribosome interaction in mycobacteria, we mixed recombinant Rsh_C666G_ and Rsh_C692F_ variants with 70S ribosomes purified from Δ*rsh* mutant of *M. smegmatis* and analyzed their occupancy on the ribosome. Ultracentrifugation on a 32% sucrose cushion was used to separate ribosome-bound from unbound Rsh. Compared to the wildtype Rsh, association with the ribosome was significantly compromised for both recombinant proteins, Rsh_C666G_ and Rsh_C692F_ (Fig. 3C). We thus conclude that Rsh constitutively interacts with the ribosome using its ZBD in non-starved cells, and this interaction is necessary for the intracellular stability of the enzyme.

**Figure 3:**
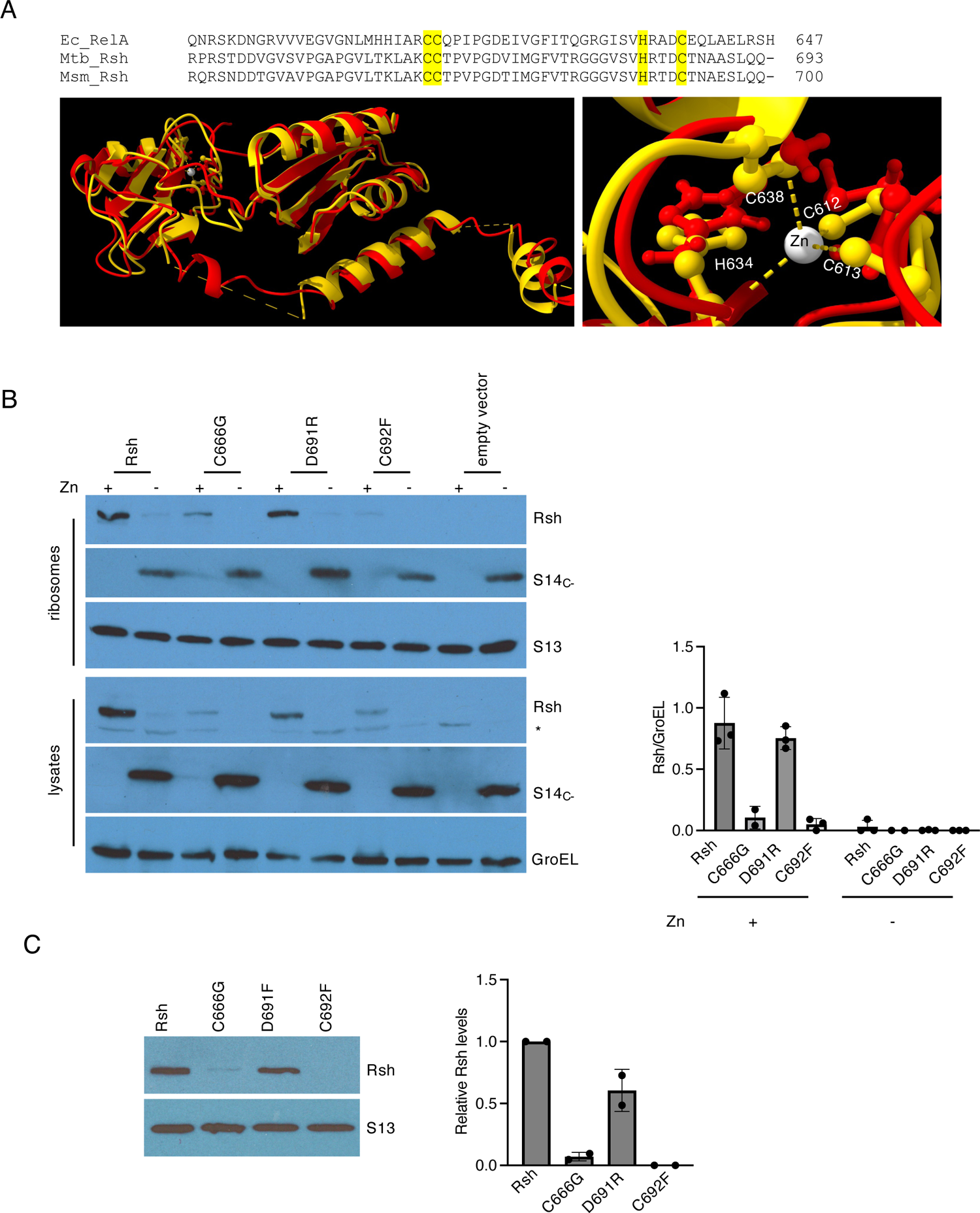
Ribosome binding residues in Rsh are critical for its intracellular stability. **A.** (Top) Sequence alignment of the C-terminal region of Rsh from *E. coli*, *M. smegmatis* and *M. tuberculosis* depicting the CCHC like zinc-finger (highlighted yellow) in the ZBD. (Bottom) Superimposition of the structure (PDBID: 5IQR) of the ZBD region from *E. coli* (yellow) and *M. smegmatis* (red). Zinc atom coordinated (shown with dashed lines) by the CCHC motif (shown as ball-and-stick model and numbered based on *E. coli* sequence) has been zoomed in the image to the right. The corresponding residues from *M. smegmatis* (shown in red ball-and-stick model) predict similarity in spatial orientation with respect to their *E. coli* counterparts. **B.** Immunoblot analysis of Rsh variants (C_666_G, D_691_R and C_692_F) in cell lysates and 70S ribosomes purified from 96-hour-old high-zinc cultures of a Δ*rsh* strain *of M. smegmatis* expressing each Rsh variant individually from an integrative plasmid (pYL238, pYL240, pYL241, pYL242 for Rsh, Rsh_C666G_, Rsh_D691R_ and Rsh_C692F_, respectively). Cultures of Δ*rsh* strain expressing WT Rsh were controls. Normalized average levels of Rsh and its variants in cell lysates from three independent experiments are shown in the plot. **C.** *In vitro* ribosome binding of recombinant Rsh and its variants. 2.4 picomoles of 70S ribosomes purified from a 96-hour old high-zinc culture of Δ*rsh* strain of *M. smegmatis* were mixed with 1.2 picomoles of Rsh or variant proteins (purified from recombinant *E. coli* harboring pYL243, pYL246, pYL247, pY248 for Rsh, Rsh_C666F_, Rsh_D691R_ and Rsh_C692F_, respectively) and the mixture was incubated for 30 minutes at 37 ^∘^C. The complex was separated by ultracentrifugation through a 32% sucrose cushion. S13 was probed as the loading control. The plot shows quantitative analysis of Rsh from two biologically independent experiments.

### Rsh interacts with the IF2 bound 70S initiation complex

We next focused on understanding the stage of translation when Rsh would most likely interact with the ribosome in a non-starved cell. We streamlined our approach based on the published structures (17–19), which delineate permissive and non-permissive translation complexes for RelA binding. While RelA can bind to a late translation-initiation intermediate (70S-fMet-tRNA_i_^Met^-mRNA with vacant A-site) (18), its binding to the ribosome during translation elongation is expected to be sterically hindered by inter-subunit rotation (17–19). We therefore hypothesized that mycobacterial Rsh would enter the translation initiation complex through one of the free ribosomal subunits and likely remain on the complex until the start of the first cycle of elongation. This hypothesis also implies that Rsh must remain associated with the ribosome during its transition from initiation to pre-elongation stages, without affecting the overall canonical conformation of the translation complex.

To investigate this possibility, we first determined the propensity of Rsh to interact with free ribosomal subunits. Purified recombinant Rsh was mixed individually with each of the two purified ribosomal subunits free of any canonical protein synthesis ligands and the macromolecules were separated on a 32% sucrose cushion. Rsh appeared to preferentially co-sediment with the LSU (Fig. 4A). We next asked if Rsh would remain in the translation complex during its transitions through the various intermediate stages of initiation, especially given the potential of a steric clash between Rsh and translation initiation factor 2 (IF2). The contact points of IF2 on the 70S initiation complex significantly overlaps with those of Rsh (17–19, 46). We therefore tested the ability of Rsh to interact with the ribosome in the presence of IF2, fMet-tRNA_i_^Met^ and mRNA *in vitro*. Dissociated ribosomes (Fig. S6) were mixed with His_6_-IF2, His_6_-Rsh, fMet-tRNA_i_^Met^, leadered mRNA, and either GTP or a non-hydrolysable analogue (GMP-PNP). The assembled 70S complex purified on 10-40% sucrose density gradient (SDG) was analyzed for the presence of IF2 and Rsh. The 70S ribosomes assembled in the presence of GTP contained Rsh but had undetectable levels of IF2 (Fig. 4B), demonstrating that Rsh can associate with the ribosome in the presence of the canonical ligands without interfering with GTP-dependent release of IF2 (47). This property of mycobacterial Rsh is in sharp contrast to *B. subtilis* Rel, which cannot form a stable complex with the ribosome without an A-site deacylated-tRNA (23). Unexpectedly, both Rsh and IF2 were detected in 70S particles assembled in the presence of GMP-PNP (Fig. 4C), suggesting that one or both proteins could be occupying the ribosome in their non-canonical states.

**Figure 4:**
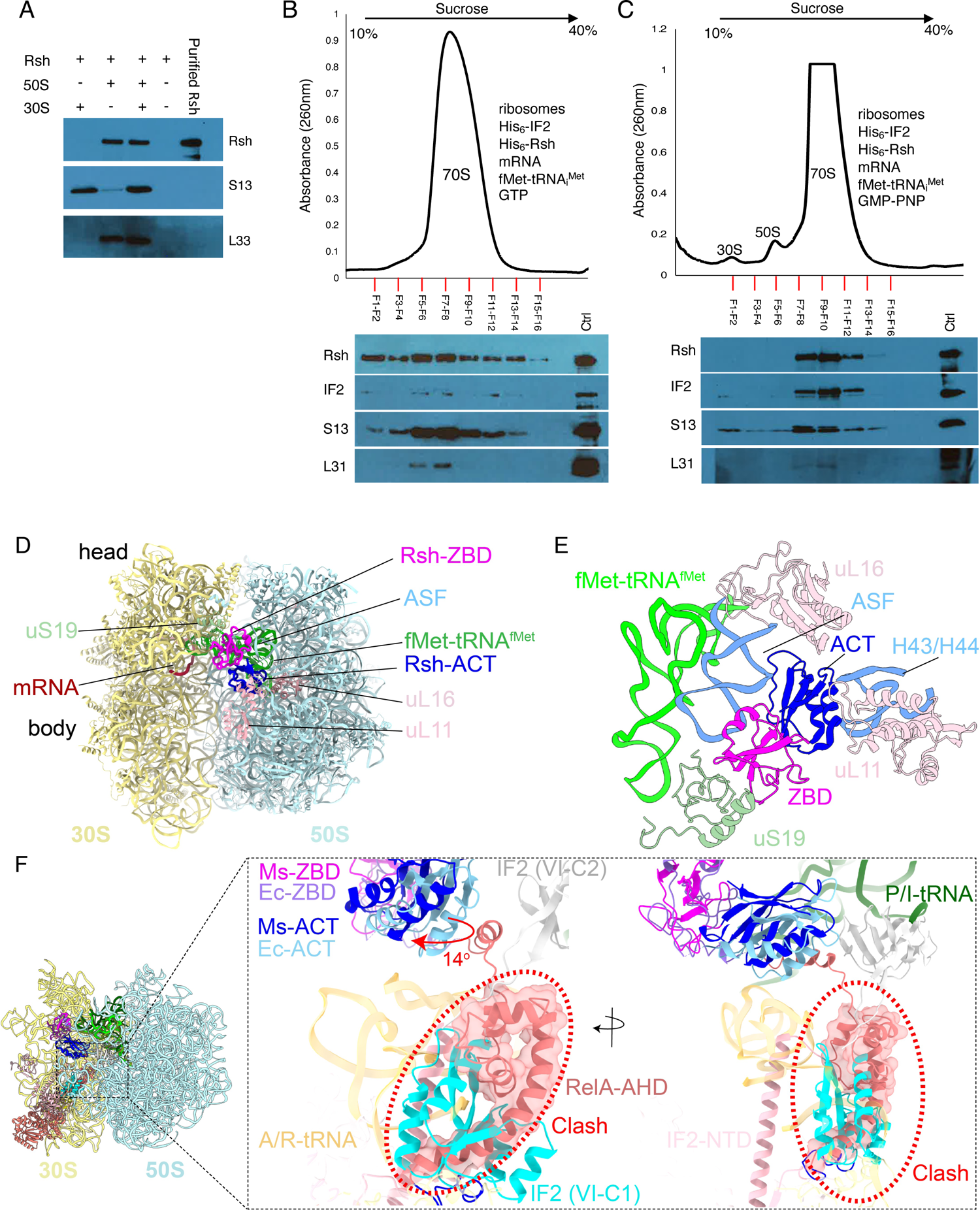
Rsh interaction with a 70S translation initiation complex. **A.** Co-sedimentation of Rsh with the 50S ribosomal subunit. Pellets resulting from ultracentrifugation on a 32% sucrose cushion were analyzed for S13 to confirm SSU, L33_C-_ to confirm LSU, and Rsh to test preferential subunit binding. Rsh in the absence of ribosomes was included as control to rule out sedimentation of free protein. Purified Rsh was loaded as marker. **B-C.** Reconstitution of Rsh-bound pre-elongation complex in the presence of GTP (B) or GMP-PNP (C). Dissociated ribosomes (Figure S6) were mixed with indicated ligands in the presence of GTP or GMP-PNP. After incubation for 15 minutes in low-magnesium (1mM MgCl_2_) buffer, recombinant Rsh was added, and magnesium was adjusted to 10 mM. The resulting mixture was incubated for additional 30 minutes. The complex was resolved in 10-40% SDG, and fractions corresponding to the 70S-peak and its margin were analyzed for Rsh, IF2 and indicated ribosomal proteins by immunoblotting using anti-Rsh, anti-His_6_ (for IF2), anti-S13 and anti-L31 antibodies. All reagents in the reactions in panels B and C were added in 5-fold molar excess relative to the ribosomal subunits. **D.** 2.7 Å resolution cryo-EM structure of the 70S-Rsh complex in late initiation state of translation, with a fMet-tRNA_i_^Met^ present in the P-site (green), mRNA (brown), and two C-terminal domains of Rsh, ZBD (magenta) and ACT (blue). The 30S and 50S subunits are shown in yellow and blue, respectively. **E**. Ribosomal components interacting with Rsh ZBD and ACT domain, along with the fMet-tRNA_f_^Met^ in the P-site (green), are shown. ZBD interacts with components of inter-subunit bridge B1a, uS19 (light green), and the A-site finger (ASF, sky blue). ACT interacts with the 50S components uL16 (sky blue), L11, and 23S rRNA helices H43/44 (sky blue). F. Superimposition of *M. smegmatis* 70S-Rsh, *E. coli* 70S-RelA (PDB ID: 5KPV) and *E. coli* 70S-IF2 (PDB ID: 6HTQ) structures, comparing their binding sites and conformations on the ribosome. Position of IF2 (VI-C1, cyan) suggests a direct steric clash (highlighted by red dotted circle) with the *E. coli* RelA’s α-helical domain (RelA-AHD, light coral). *M. smegmatis* Rsh-ACT (Ms-ACT, blue) is rotated by ∼14° (indicated by a red arrow) relative to the *E. coli* RelA-ACT (Ec-ACT, light blue), while the ZBD domains (Ms-ZBD, magenta and Ec-ZBD in light purple) have the same position.

### Cryo-EM structure reveals unknown ribosome-Rsh interactions

To gain further insight into the Rsh-bound *M. smegmatis* 70S ribosome in a translation initiation complex, we performed single-particle cryo-EM to determine the structure of the 70S ribosome assembled in the presence of IF2, Rsh, fMet-tRNA_i_^Met^, leadered mRNA, and GMP-PNP. 3D classification of the cryo-EM dataset (Figs. S7, S8, Table S3) revealed a unique class of Rsh-bound ribosome in a classical non-rotated state, but none of the classes showed any density for IF2. The Rsh-bound 3D class refined to 2.7 Å resolution showed densities only for two C-terminal domains of Rsh, ZBD and ACT, along with a fMet-tRNA_i_^fMet^ bound at the ribosomal P site, while all other domains were found to be disordered (Figs. 4D, 4E, S7 and S9). The ZBD interacts with both the SSU and LSU: specifically, protein uS19 of the SSU and 23S rRNA helix H38 of the LSU (also referred to as ASF) (Fig. 4E). These interactions resemble those reported in the structures of *E. coli* RelA and *B. subtilis* Rsh on the ribosome (18, 20). The Rsh-ACT domain occupies the inter-subunit space and interacts with LSU proteins uL16, uL11, and the 23S rRNA helices H43-44 at the base of the protein L7/L12 stalk (Fig. 4E).

Previous structural studies suggest IF2 and RelA binding to the ribosome would be mutually exclusive (18, 46). However, our data shows co-existence of the two proteins in a reconstituted 70S translation initiation complex (Fig. 4C). To begin to resolve this issue, we superimposed either our mycobacterial structure or the *E. coli* 70S ribosome-RelA structure (18) with 70S-IF2-GMP-PNP initiation complexes (46). Indeed, RelA’s alpha-helical domain (AHD) would directly clash with domain VI-C1 of IF2 (Fig. 4F). In tertiary structure, the AHD of *E. coli* RelA lies adjacent to the ACT domain while the ZBD is anchored to the ribosome via its interactions with uS19 and ASF (18). In our structure the rotational angle between the ACT and ZBD domains was ∼14 degrees greater than that in *E. coli* RelA (Fig. 4F) and AHD was disordered, presumably displaced from its canonical position, thus avoiding a clash with domain VI-C1 of IF2 (Fig. 4F). The ACT domain is known to exhibit some conformational variations with respect to the ZBD domain in the previously published structures (18, 20), but an additional rotation by 14 degrees is unprecedented. It is possible that the adopted conformation of the ACT domain in our structure is to accommodate the exiting IF2 molecule upon completion of the translation initiation phase.

Compared to the *E. coli* RelA and *B. subtilis* Rel, mycobacterial Rsh shows an overall similarity in tertiary structures of ZBD and ACT domains, although the rotation of the ACT domain translates into its shift by ∼6 Å towards the LSU’s L7/L12 stalk base (Fig. 5A, B). This altered conformation of ACT in Rsh is stabilized through extensive interactions with the uL11 and the 23S rRNA helices H43/44 (Fig. 5C). Specifically, uL11 lysines, 89 and 96 interact with Y791, D789 and H776 of Rsh-ACT. Further, Rsh-ACT S779, N783, D789 and Y788 interact with the 23S rRNA helices H43-44 phosphate backbones. These interactions are in sharp contrast to the *E. coli* ribosome-RelA structures, where uL11 interacts with the A/R tRNA but not directly with RelA (18). Thus, our structure reveals a unique conformational state of the ribosome-bound Rsh. Multiple sequence alignment of the amino acid sequences does not reveal any apparent substitutions, insertions, or deletions that can explain the altered orientation of the ACT domain in *M. smegmatis* Rsh (Fig. S10). Given that our structure was obtained in the presence of IF2-GMP-PNP, a key difference from the *E. coli* ribosome-RelA complex, a possible role of IF2 in inducing the rotation of ACT cannot be ruled out.

**Figure 5:**
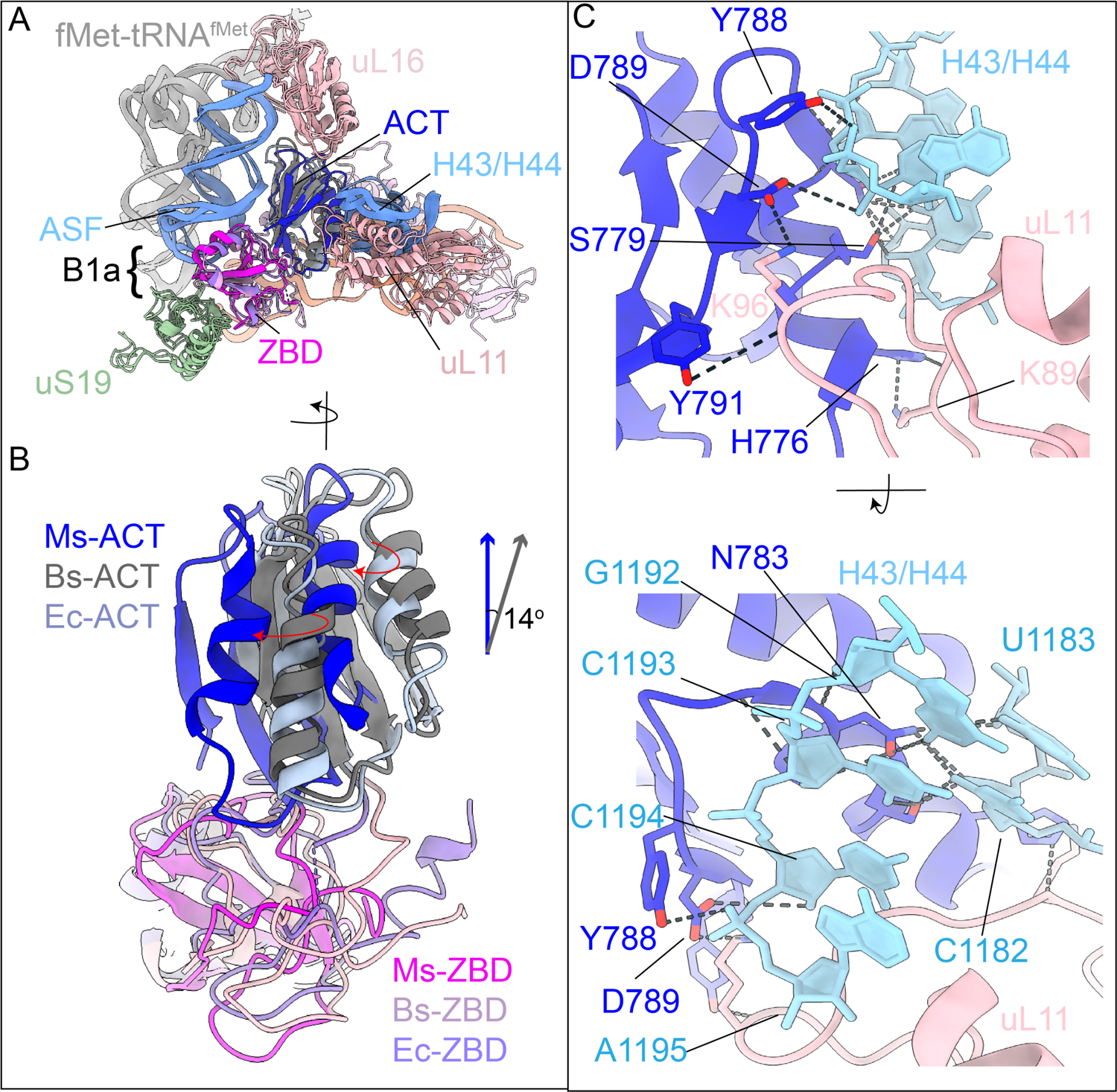
The Rsh-ACT domain interacts with the L7/L12 stalk base. **A.** Superimposition of the Rsh/Rel A interacting components from the 70S-Rsh/RelA complex structures from *M. smegmatis* (Ms) 70S-Rsh (ZBD, magenta; ACT, blue), *B. subtilis* (Bs) 70S-Rsh (ZBD, light pink; ACT, gray), and *E. coli* (Ec) 70S-RelA (ZBD, light purple; ACT, steel blue). Only components interacting with RelA/Rsh, L7/L12 stalk base (H43/H44 in blue), A-site finger (ASF in blue) uL16 and uL11 (light pink), the A/R tRNA (light coral) and uS19 (sea green), along with the P-site tRNA (light gray), are displayed. **B.** Superimposition of the C-terminal ZBD and ACT domain from Ms, Bs, and Ec (color-coded as in panel A) highlighting the rotation in Ms-ACT domain with respect to Ms-ZBD. **C.** Interactions of Ms Rsh-ACT (blue) with L11 (light pink) and the 23S rRNA helices H43/44 (light blue) are shown from two viewing directions.

Interactions stabilizing Rsh in mycobacterial structure are primarily through LSU’s ASF, uL16, uL11, and rRNA helices H43/44, suggesting that Rsh associates primarily with the LSU, corroborating the results of our binding experiments that show association of Rsh with free LSU. The rRNA helices H43/44 and uL11 together constitute the L7/L12-stalk base of LSU and are known to be mobile during elongation (48) and recycling (49) phases of translation. We thus propose that intensive contacts between the L7/L12-stalk base and the Rsh-ACT domain would transiently lock the ribosome in a stalled state before the start of the translation elongation cycle, as a mechanism to survey the aminoacylation state of the incoming tRNA to the A site. A direct contact of Rsh with uL11 is distinct from *E. coli* RelA, which binds to the ribosome in uL11-independent manner, even though uL11 and a deacylated-tRNA at the A-site are required for (p)ppGpp synthesis (21, 50–52). While the structural changes in *M. smegmatis* complex here could be induced by the presence of IF2, we cannot exclude the possibility that these are intrinsic *to M. smegmatis* Rsh.

The superimposed structures also reveal some species-specific differences in Rsh/RelA/Rel, which could probably explain differences in their ribosome binding affinities (Fig. S10). The ZBD domains interact with uS19 and ASF in all three cases, however, compared to *M. smegmatis*, *B. subtilis* Rel has additional interactions — two with uS19, one with ASF, and one with uS13 (Fig. S10). The loss of the additional interactions with uS19 and ASF in *M. smegmatis* is likely due to substitution at corresponding positions with non-polar amino acids (Fig. S10). However, *M. smegmatis* Rsh-ZBD could potentially interact with uS13 since it has an Arg (R681) corresponding to K617 in *B. subtilis*. The *E. coli* RelA ZBD also has two additional interactions compared to *M. smegmatis*, one with uS19 and the other with ASF (Fig. S10). However, *E. coli* RelA ZBD has a Gln (Q627) corresponding to K617 and R681 in *B. subtilis* and *M. smegmatis*, respectively, which might not interact with uS13 due to its shorter side chain. Since the density corresponding to ZBD is relatively strong in our structure, we designate it as the primary anchoring domain of mycobacterial Rsh.

### Exclusion of Rsh from Mpy-bound ribosomes

The requirement of the ribosomes for the intracellular stability of Rsh, combined with its loss in cells harboring Mrf and Mpy-bound hibernating ribosomes, suggests that Rsh is not able to bind to hibernating ribosomes. Mpy and Rsh bind at two physically distinct sites on the unrotated SSU of the ribosome (Fig. 6A). The Mpy N-terminal domain binds to the SSU spanning the decoding center by interacting extensively with the 16S rRNA components that converge from platform, body, and head regions at the SSU’s neck (38). By contrast, the Rsh ZBD interacts near the inter-subunit bridge B1a (53) forming components, including uS19 of the SSU and the LSU’s ASF (54) (Figs. 4E and 6A). Since the binding sites of the two factors do not overlap, we superimposed the structures of Rsh- and Mpy-bound *M. smegmatis* 70S ribosomes to understand the accessibility of Mpy-bound hibernating ribosome to Rsh binding. Both the ribosomes are in an unrotated state, but the SSU head domain in the Rsh-bound structure is tilted more towards the LSU in comparison to the Mpy-bound ribosome structure (Fig. 6B). We predict that the untilted SSU head in the MPY-bound state will prevent stable interactions between Rsh-ZBD and uS19 (Fig. 6C). ZBD is the primary anchoring domain of Rsh (18, 20), thus, any alteration in the bridge B1a configuration in the Mpy-bound state is likely to directly inhibit Rsh binding, thereby rendering ribosome-free labile Rsh. Thus, reduction in the intracellular Rsh during ribosome hibernation could be a combined outcome of both a decrease in the synthesis of new Rsh and the occlusion of pre-existing Rsh from the hibernating ribosome.

**Figure 6:**
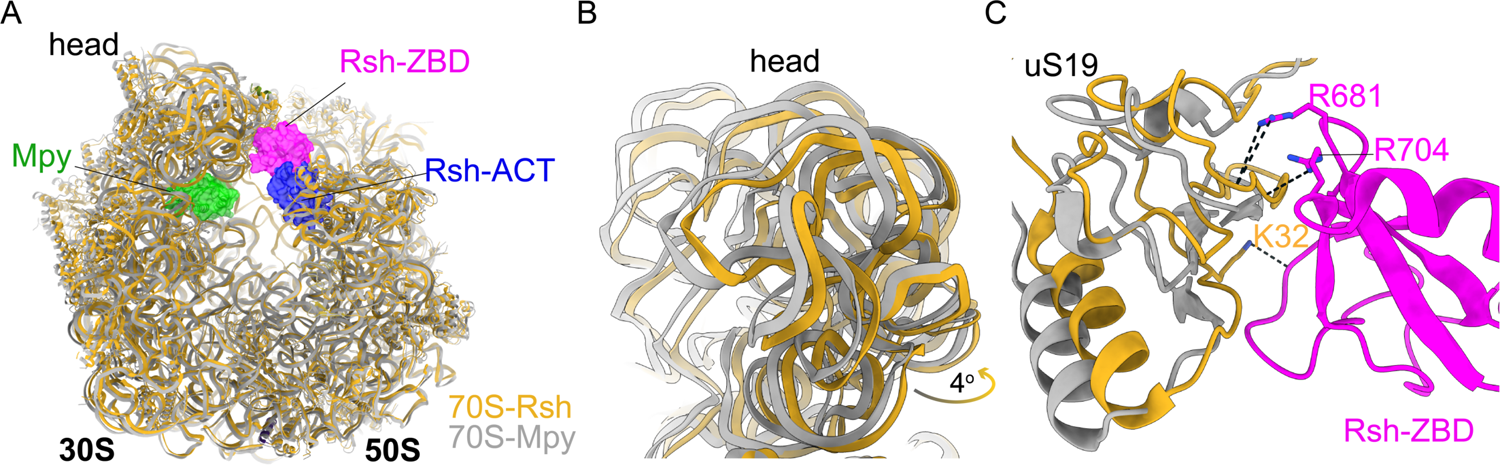
Structural basis of Rsh exclusion from a Mpy-bound 70S ribosome. **A**. Superimposition of Ms 70S-Mpy (gray, PDB ID:6DZI) and Ms 70S-Rsh (gold) structures. Rsh (ZBD, magenta; ACT, blue) and Mpy (green) are shown in the space-filled rendering. **B**. The 16S rRNAs corresponding to head domain of the 30S subunit in panel A has been enlarged to highlight their relative position in the two functional states. **C**. The position of uS19 (gray) in the Mpy-bound state would disrupt the observed interactions between Rsh-ZBD (magenta) and uS19 (orange).

## Discussion

The stringent response, characterized by Rsh-dependent synthesis of (p)ppGpp and subsequent transcriptional reprogramming by the alarmone, is among the central adaptation mechanisms in nutrient-starved mycobacteria. Yet, a significant knowledge gap exists in the mechanism underlying starvation sensing by Rsh. Structural studies with *E. coli* RelA provide mechanistic details on how Rsh is activated by the presence of deacylated-tRNA at the A-site in a stalled translation-elongation complex. However, it remains unclear how Rsh would encounter such a complex *in vivo*. Two models have emerged from recent studies: a) the hopping model in which a Rel-deacylated-tRNA complex is formed prior to its entry at the A-site of the elongation complex (22), and b) starvation-induced formation of a weak RelA-ribosome complex prior to its stabilization by deacylated-tRNA (23). Both models however preclude RelA-ribosome interaction in non-starved cells, which is inconsistent with the evidence of constitutive RelA-ribosome interaction throughout the growth of *E. coli* and *P. aeruginosa* cells (33, 55). Here, we propose a new model of Rsh recognition of a stalled translation complex in mycobacteria. In this model, which we call the surveillance model, Rsh constitutively interacts with ribosomes during the formation of the initiation complex, and its transition to the pre-elongation stage (Fig. 7A). Consequently, entry of an amino acylated-tRNA would allow elongation, accompanied by inter-subunit rotation that would displace Rsh from the complex (Fig. 7B) (17–19). By contrast, entry of a deacylated tRNA into the A-site would trigger Rsh activation and (p)ppGpp synthesis (Fig. 7D). Free Rsh appears to be labile (Fig. 7G). To maintain steady state levels Rsh must therefore associate with free LSU entering the translation cycle (Fig. 7C), or another stalled elongation complex with deacylated-tRNA at the A-site (Fig. 7E). The latter interaction is expected to be uncertain and less frequent given the low abundance of stalled elongation complexes relative to large abundance of free LSU. Given that Rsh abundance in an actively growing cell is ∼200-fold less compared to the ribosome (21), a rapid rate of translation would likely maintain a sufficiently large pool of free LSU transitioning between termination and initiation cycles, thereby ensuring a stable population of Rsh-ribosome complex. The predictable fates of Rsh in the surveillance model renders the model more stringent than either the original or the refined hopping model, both of which rely on random encounter of Rsh with a deacylated-tRNA and/or a stalled ribosome.

**Figure 7:**
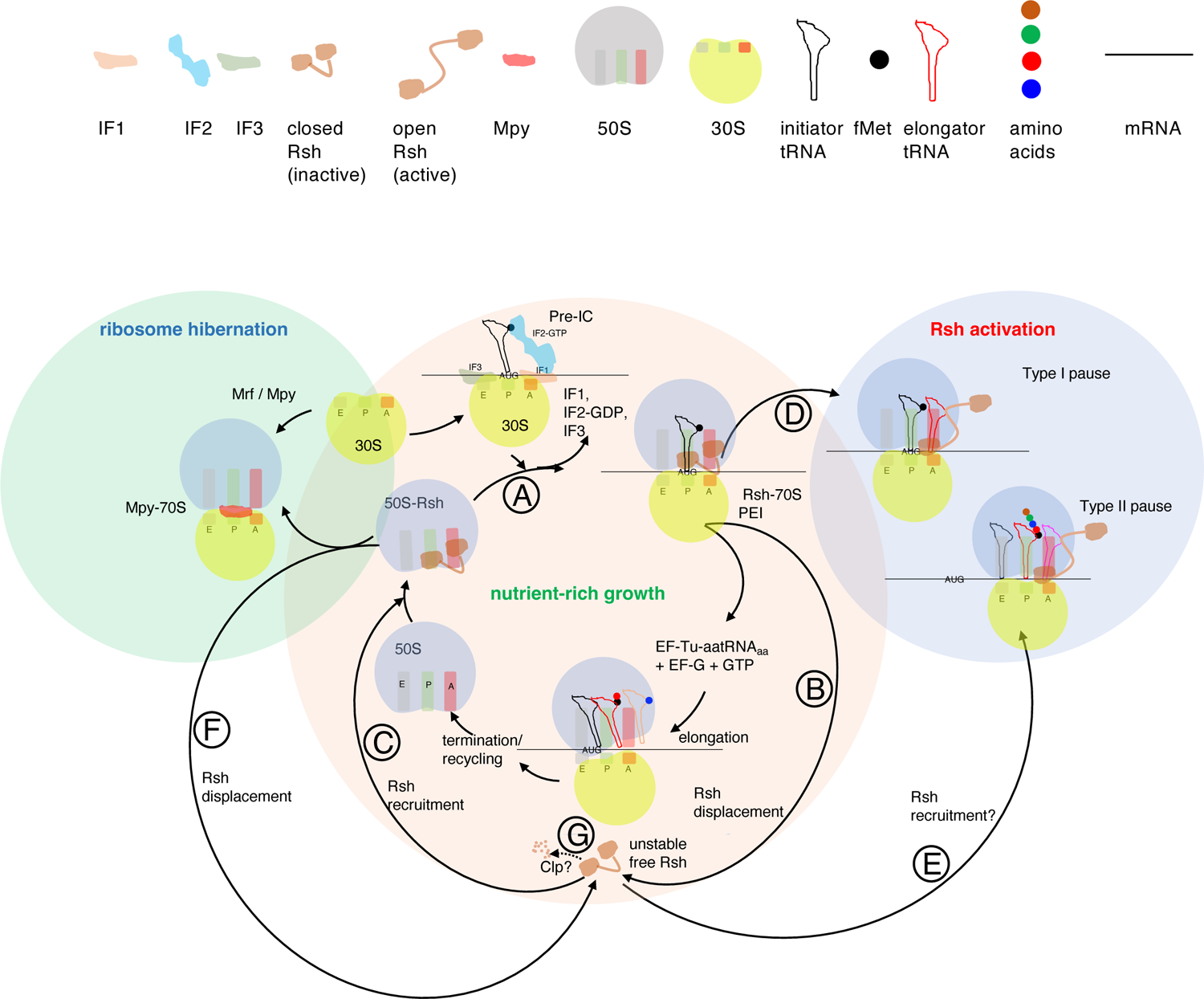
Model of a dynamic interaction of Rsh with the ribosome in mycobacteria, influenced by three distinct conditions. In a nutrient-rich condition supporting high translation activity, labile Rsh predominantly remains associated with the free 50S subunit, which confers stability to the protein. Rsh-50S complex joins with a 30S pre-initiation complex (pre-IC). Following the release of initiation factors, Rsh likely remains in the pre-elongation intermediate (PEI), which in an unrotated conformation is ready to elongate upon entry of the first A-site tRNA (step A). Rsh is likely sterically displaced during inter-subunit rotation associated with the first cycle of elongation (step B). Free Rsh associates with 50S subunit (step C) and the cycle continues until a deacylated tRNA enters the A-site at the second codon (type I pause), which leads to activation of Rsh (step D). Rsh could also potentially be activated if free Rsh interacts with a stalled ribosome downstream of the second codon (Type II pause) (step E), although its frequency could be limited by relatively low abundance of such stalled complexes compared to the free 50S subunit. Conditions inducing hibernating ribosomes, which are likely less accessible to Rsh, would sequester ribosomes from the translation cycle and displace Rsh (step F), thereby rendering Rsh labile in a Clp protease dependent manner (step G).

The lack of Rsh under conditions such as those inducing ribosome hibernation further supports the model that optimum protein synthesis and accessibility to LSU entering the translation cycle is critical for the intracellular stability of Rsh (Fig. 7F). This loss of Rsh further differentiates the physiological conditions for ribosome hibernation and the stringent response. The stringent response is an upstream response before ribosome hibernation; a logical order given that the stringent response reprograms gene expression to fine-tune metabolism, while ribosome hibernation directly shuts down global protein synthesis. Moreover, restoration of (p)ppGpp synthesis in a zinc-depleted Δ*mrf* strain implies that Rsh-ribosome interaction and Rsh activation are relatively less sensitive to intracellular depletion of free zinc. We hypothesize that the coordinating zinc residues of Rsh have high affinity for zinc, supporting continued interaction with the ribosome even during the declining intracellular level of free zinc. This would allow activation of Rsh under low-zinc conditions before the induction of Mpy-dependent ribosome hibernation. Zinc is an essential nutrient that binds to 6-10% of all proteins in bacteria (56). Notably, several amino acyl tRNA synthetases are zinc binding proteins (56). A lack of zinc in the active site of threonyl-tRNA synthetase inactivates the aminoacylation activity of the enzyme (57).

Moreover, after alanine (26%) threonine (17%) is the second-most frequent amino acid to follow the initiator Met in the top 183 most translated genes in *M. smegmatis*. Limitation in zinc therefore will directly increase the level of deacylated-Thr-tRNA, activating the stringent response through the surveillance model. Thus, the intracellular availability of zinc in mycobacteria could potentially be a shared signal for the sequential onset of the stringent response and ribosome hibernation, mediated directly through sensing the translation activity of the ribosome. Beyond zinc starvation as the documented condition leading to mycobacterial ribosome hibernation, it is likely that other conditions inducing ribosome hibernation would also attenuate the stringent response in a similar manner.

The Clp-dependent loss of ribosome-free intracellular Rsh, while unveiling a novel mechanism of regulation of Rsh activity *in vivo*, does not entirely rule out the CTD-dependent autoinhibition of the synthase activity through oligomerization: a model primarily proposed from *in vitro* characterization of the enzyme (24–26). However, the evidence of ribosome-bound Rsh as the dominant species *in vivo* suggests that Rsh oligomers, if any, would be a relatively minor molecular species in the cell. The predominance of ribosome-bound Rsh *in vivo* further suggests a reversible transition between the synthase and the hydrolase activities may occur through differential allosteric effects of ppGpp and GDP/ATP binding to the hydrolase and the synthase domain, respectively, as shown by Tamman et al. (28).

In summary, we propose a unique structure-based mechanism of mycobacterial Rsh recognition of translating ribosomes stalled with deacylated A-site tRNA, in which the enzyme predominantly surveys the amino acylation status of the tRNA during the first cycle of elongation to activate the stringent response. This mechanism of Rsh activation is programmed to slow down upon induction of ribosome hibernation. These findings will serve as a new molecular frame of reference for further investigation of the relationship between the stringent response and ribosome hibernation in mycobacteria.

## Materials and Methods

### Bacterial growth medium

*Mycobacterium smegmatis* (mc^2^155) and its variant strains were grown shaking at 200 rpm at 37°C in Middlebrook 7H9 with ADC enrichment (5% albumin, 2% dextrose, 0.85% sodium chloride and 0.003% catalase) and 0.05% (v/v) Tween80. For ribosome purification, cells from saturated cultures were washed with PBS + 0.05% (v/v) Tween80 for three times and then inoculated at 1:100 dilution in Sauton’s medium with either 1 mM ZnSO_4_ for high-zinc condition or 1 μM zinc-chelator (TPEN, N,N,N’,N’-Tetrakis(2-pyridylmethyl)ethylenediamine) for low-zinc condition. 7H10ADC agar plate was used for selection of recombinants and colony growth (*M. smegmatis*). *Escherichia coli* (GC5 or BL21) was grown in LB broth or LB agar at 37 °C. Zeocin (25 μg/mL), kanamycin (20 μg/mL), hygromycin (150 μg/mL for *M. smegmatis* and *E. coli*, carbenicillin (50 μg/mL), and apramycin (5 μg/mL) were used for selection as necessary. For recombinant strain harboring acetamide-inducible promoter, cells were cultured in high- or low-zinc Sauton’s medium with 0.2% sodium succinate and induced with 0.2% acetamide.

### Construction of recombinant plasmids and strains

A list of plasmids, bacterial strains used in this study is provided in Table S1. A list of oligonucleotides used in this study is provided in Table S2. For construction of Δ*rsh* in *M. smegmatis*, the gene was replaced with a zeocin-resistance marker using a PCR-based modified version of the recombineering strategy as previously described (38). For creating C_666_G, D_691_R and C_692_F variants of Rsh, the Rsh ORF with 500 bp of upstream sequence was PCR-amplified and cloned between KpnI and XbaI sites of the integrative vector pMH94 (39), resulting in the recombinant plasmid pYL238. Mutations at the respective sites were introduced using pYL238 as a template by PCR-based mutagenesis using the primer pairs listed in Table S2, resulting in pYL240 (C_666_G), pYL241 (D_691_R) and pYL242 (C_692_F). Mutations were sequence-confirmed and the plasmids pYL238, pYL240, pYL241 and pYL242 were electroporated into the mc^2^155:Δ*rsh* strain of *M. smegmatis*. Cells expressing the mutant variants were cultured in nutrient-rich medium and analyzed for Rsh levels by immunoblotting. The integrative plasmid with anhydrotetracycline(ATc)-inducible CRISPRi-dCas9 system targeting the *clpP1* expression was constructed and donated by Dr. Keith Derbyshire as a part of the Mycobacterium Systems Resources (MSR)(58). The *M. smegmatis* strain (YL9) harboring the plasmid was cultured in high- and low-zinc and ClpP1 depletion was induced by ATc as described previously (39).

### Recombinant protein expression and purification

Rsh from *M. smegmatis* (Msmeg_2965) and its variants (Rsh_C666G_, Rsh_D691R_ and Rsh_C692F_) with an N-terminal His tag fusion were subcloned in pET21b vector using NdeI and HindIII. The plasmids were introduced into BL21 (DE3) pLysS cells for Rsh expression and purification. Briefly, the cells were grown in LB medium to log phase (OD_600_ of 0.7) at 37 °C, when RSH expression was induced with 0.5 mM IPTG for 4 hours at 37 °C. Induced cells were harvested (8000 rpm, 20 minutes at 4°C in Thermo Scientific™ Fiberlite™ F12-6 × 500 Fixed-Angle Rotor) and then resuspended in N-I buffer (50 mM Tris (pH 8.0), 300mM NaCl, 5% glycerol, 10 mM imidazole, 1mM PMSF). The cells were further sonicated (Amplitude 30%, 6 cycles of 10 sec on and 60 sec off for) at 4 °C and the soluble protein in the supernatant was separated from the debris by centrifugation (13000 rpm, 20 minutes, 4 °C in Thermo Scientific™ Fiberlite™ F12-6 × 500 Fixed-Angle Rotor) and incubated with pre-equilibrated Ni-NTA resin for 1 hour at 4 °C. After incubation, Ni-NTA resin was washed three times with N-II buffer [50 mM Tris (pH 8.0), 300mM NaCl, 5% (v/v) glycerol, 30 mM imidazole]. Rsh was then eluted with stepwise increments of imidazole (from 60 mM, to 100mM and to 250 mM) in a buffer containing 50 mM Tris (pH 8.0), 300mM NaCl and 5% (v/v) glycerol. Peak fractions with were pooled and imidazole was removed by dialysis in the buffer containing 50 mM Tris (pH 8.0), 300mM NaCl and 30% (v/v) glycerol. For recombinant IF2 (MSMEG_2628) with N-terminal His_6x_ tag, the full-length codon optimized gene with N-terminal 6x-His-sequnce and NdeI-XhoI sites was synthesized (IDT DNA Inc.) (Table S2). The synthetic fragment was cloned into a pET21a vector. Recombinant protein was expressed in BL21-DE3 (pLysS) and purified on Ni-NTA matrix as detailed for His_6_-Rsh above.

### Ribosome purification

Rsh bound ribosomes were purified with a few modifications from the previously described protocol (38). *M. smegmatis* cells were grown for the specified time in 500 mL of Sauton’s media containing 0.05% Tween 80 and either 1 mM ZnSO_4_ or 1 μM TPEN. Cells were harvested (8000 rpm for 20 minutes in a Thermo Scientific™ Fiberlite™ F12-6 × 500 Fixed-Angle Rotor) and pulverized 6 times at 15 Hz for 3 minutes in a mixer mill (Retsch MM400). The lysates were mixed with 20 mL of HMA-10 buffer (20 mM HEPES-K pH7.5, 30 mM NH_4_Cl, 10 mM MgCl_2_, 5 mM β-mercaptoethanol) and centrifuged for 30 minutes at 30,000g in Thermo Scientific™ Fiberlite™ F12-8 × 50 Fixed-Angle Rotor. The supernatants were collected in the Beckman PC ultracentrifuge tubes and further centrifuged for 2 hours and 15 minutes at 42800 rpm in a Beckman rotor Type 70Ti. Pellets were soaked in 4 mL of HMA-10 buffer in an ice water bath overnight, and then homogenized for 30 minutes. The homogenized pellets were treated with 3 units/mL Rnase-free Dnase (Ambion) for two hours at 4 °C. The contents were transferred to Beckman PA tubes and centrifuged at 22,000 g for 15 minutes in Thermo Scientific™ Fiberlite™ F12-8 × 50 Fixed-Angle Rotor. The supernatants were collected into Beckman ultracentrifuge PC tubes and centrifuged for 2 hours and 15 minutes at 42,800 rpm in a Beckman rotor Type 70Ti. The pellets were dissolved in 1 mL HMA-10 buffer and centrifuged for 10 minutes at 10,000 g. The supernatants containing the crude ribosomes were collected and quantified by measuring their optical density at 260 nm. The crude ribosome preparations were then layered on top of 37 mL sucrose density gradients (10%-40%) and centrifuged for 16 hours at 24,000 rpm in a Beckman rotor SW 28. The 70S ribosome fractions were collected after fractionating the sucrose gradient in a 260 nm Brandel gradient analyzer as previously described (59). The pooled 70S ribosome fractions pelleted by ultracentrifugation at 42,800 rpm for 3 hours in a Beckman rotor Type 70Ti were suspended in HMA-10 buffer (20 mM HEPES-K pH7.5, 30 mM NH_4_Cl, 10 mM MgCl_2_, 5 mM β-mercaptoethanol) and quantified by measuring absorbance at 260 nm.

MPY-bound ribosomes and other high-salt washed ribosomes were purified exactly as described previously (38, 39). The steps up to homogenization were same as described above. After homogenization, 4 mL of the homogenized suspension was treated with 3 units/mL Rnase-free Dnase (Ambion) for one hour at 4 °C, then 4 mL of HMA-0.06 buffer (20 mM HEPES-K pH 7.5, 600 mM NH_4_Cl, 10 mM MgCl_2_, 5 mM β-mercaptoethanol) was added to the suspension, which was further incubated at 4 °C for 2 hours. The content was transferred to Beckman PA tubes and centrifuged at 22,000 g for 15 minutes in Thermo Scientific™ Fiberlite™ F12-8 × 50 Fixed-Angle Rotor. Supernatants were collected and layered on top of 8 mL 32% sucrose solution in HMA-10 buffer and centrifuged for 16 hours at 37000 rpm in a Beckman rotor Type 70Ti. The supernatant was discarded, and the pellet was rinsed with HMA-10 buffer to remove the brownish material. The crude ribosome was resuspended in 1 mL HMA-10 buffer and centrifuged for 10 minutes at 10,000 g. Supernatant containing the crude ribosome was collected and quantified by measuring optical density at 260nm. 70S ribosome from the crude ribosome was further separated on a 10-40% sucrose density gradient and fractionated as indicated above.

### In vitro reconstitution of Rsh with 70S initiation complex

#### Co-sedimentation of Rsh with the ribosomal subunits

70S ribosomes were purified from low-zinc culture of isogenic Δ*rsh*/Δ*mpy*/Δ*mrf* triple mutant of *M. smegmatis*. Subunits were dissociated from the purified 70S ribosomes by incubation at 37°C for thirty minutes in HMA-1 buffer (20 mM HEPES-K pH7.5, 30 mM NH_4_Cl, 1 mM MgCl_2_, 5 mM β-mercaptoethanol. The resulting subunits were then purified from a 10-40% sucrose density gradient. Individual subunits equivalent to A_260_ of 0.3 (7.2 pmol) were mixed with 1.8 pmol of purified recombinant Rsh in 100 μL binding buffer (20 mM HEPES-K pH7.5, 100 mM NH4Cl, 10 mM MgCl_2_, 5 mM β-mercaptoethanol, 50mM KCl), and the mixture was incubated for thirty minutes at 37°C. The reactions were then layered on 100 μL of 32% sucrose cushion and centrifuged at 42,800 rpm for 3 hours in a Beckman TLA 100 rotor using the Optima-Max TL ultracentrifuge (Beckman Coulter). After centrifugation, the supernatant was discarded, and the ribosome pellet was resuspended in 20 μL HMA-10 buffer. The abundance of Rsh was further determined by immunoblotting using the protocol described below Rsh, and S13, and L33.

#### Preparation of fMet-tRNA_i_^Met^

An *E. coli* culture containing a plasmid borne expression vector for tRNA_i_^Met^ under control of the LacI repressor was grown to an optical density at 600 nm of 0.4-0.6 at which time IPTG was added to induce expression of the tRNA. After 8 h the cells were harvested, lysed by phenol, and total tRNA isolated using a series of differential precipitation steps (60). Approximately half of the recovered tRNA was a substrate for charging by MetRS. The charging reaction (2 mL) contained approximately 200 nmoles of total tRNA and 10 nmoles of MetRS in the presence of 1 mM methionine, 2 mM ATP, 10 mM MgCl_2_, 20 mM KCl, 4 mM DTT, 50 μg/mL BSA, and 50 mM Tris-HCl pH 7.5. After 10 min at 37 °C a 4.0 μL aliquot of the reaction was removed for determination of the charging efficiency as described by Gamper et *al* (61). The remaining reaction was incubated for 10 min at 37 °C with 1.7 µmoles of 10-formyltetrahydrofolate and 20 nmoles of methionyl formyl transferase to convert Met-tRNA_i_^Met^ to fMet-tRNA_i_^Met^. The reaction was quenched by adding a 0.1 volume of 2.5 M NaOAc pH 5.0. Following an equal volume pH 5 phenol-chloroform-isoamyl alcohol extraction (80:17:3), the tRNA was ethanol precipitated, dissolved in 300 µL 25 mM NaOAc pH 5.0, and stored at −70 °C. Approximately 200 nmoles of tRNA was recovered, of which 45% was fMet-tRNA_i_^Met^.

#### Reconstitution of Rsh-bound 70S initiation complex

*70S* ribosomes prepared from a low-zinc culture of isogenic Δ*rsh*/Δ*mpy*/Δ*mrf* triple mutant of *M. smegmatis* were dialyzed at 4°C with low-magnesium buffer (20 mM HEPES-K pH7.5, 30 mM NH_4_Cl, 0.5 mM MgCl_2_, 5 mM β-mercaptoethanol) to dissociate the 30S and 50S subunits. Two rounds of dialysis were performed, with fresh buffer change every four hours. Subunit dissociation was confirmed by loading 2.4 picomoles of the dialyzed ribosomes onto a 10-40% sucrose density gradient.

Another 2.4 pmoles of dissociated ribosomes were mixed with five-fold molar excess of fMet-tRNA_i_^fMet^, a synthetic mRNA template (GGCAAGGAGGUAAAAAUGUUCAAAAAA-Flour) (IDT, USA), recombinant His_6_-IF2, recombinant His_6_-Rsh, and either GTP or GMP-PNP (Sigma-Aldrich) in a 100 μL reaction in buffer I (20 mM HEPES-K pH 7.5, 30 mM NH_4_Cl, 1 mM MgCl_2_, 5 mM β-mercaptoethanol), and the mixture was incubated at 37 °C for 15 minutes to allow formation of the initiation complex on the 30S subunit. Then 25 μL of buffer II (20 mM HEPES-K pH7.5, 30 mM NH4Cl, 50 mM MgCl2, 5 mM β-mercaptoethanol, 600mM KCl) was added to the mixture, thereby bringing the final concentration of MgCl_2_ and KCl to 10mM and 120 mM, respectively. The reaction was then incubated for an additional 30 minutes at 37°C to allow joining of the 50S subunit. Resulting complexes were either used for structural analysis or resolved on a 15 mL 10-40% sucrose density gradient centrifuged at 35000 rpm for 135 minutes using SW41 rotor on an Optima L-90K ultracentrifuge (Beckman Coulter), and the resolved particles were fractionated as previously described (59). For immunoblotting analysis, content of the indicated fractions were precipitated using Methanol-chloroform extraction as previously described (39). Briefly, each fraction was vortexed with 4X volume of methanol for 5-10 seconds and centrifuged for 10 seconds at 13,200 rpm on a benchtop centrifuge before adding one volume of chloroform. The mixture was further centrifuged at 13,200 rpm for 60 seconds, after which 3X volume of water was added and the mixture was vortexed before further centrifugation at 13,200 rpm for 60 seconds. The upper aqueous layer was discarded while retaining the interface and the organic layer, which was mixed with 3X volume of methanol, vortexed and centrifuged at 13,200 rpm for 2 minutes. The supernatant was discarded, and the resulting pellet was dried and resuspended in 0.1 M Tris containing 8M urea for further analysis of proteins by immunoblotting.

### Immunoblotting

Total cell lysates (10 μg) or purified 70S ribosomes (2.4 pmoles) were used for detecting S13, MPY, Mrf-FLAG, and S14_C-_, while 24 pmoles ribosome were used for detecting Rsh. Samples were resolved on 8% SDS-PAGE (for FLAG, Rsh and GroEL analysis) or 12% SDS-PAGE (for MPY, S14_C-_ and S13), and proteins were probed with anti-FLAG (1:5000, Genescript), anti-GroEL (1:5000, Enzo), anti-RelA (1:5000), anti-S13 of *E. coli* (1:100, The Developmental Studies Hybridoma Bank), endogenously raised anti-MPY (1:5000) and anti-S14_C-_ (1:2000) antibodies.

### ppGpp extraction and analysis

Cells of specified *M. smegmatis* strains were grown in Sauton’s medium with either 1 mM Zn or 1 μM TPEN for 96 hours, after which 1 mL of the culture was labelled with 100 DCi/mL of ^32^P KH_2_PO_4_ (Perkin Elmer, 900-1100mCi/mmol) for 6 hours on a shaker at 37 °C. Labeled cells were washed once with TBST (20mM Tris pH7.6 + 150 mM NaCl + 0.05% Tween 80). Half of the cells (0.5 mL) were pelleted and resuspended in 25 μL of 4M formic acid and kept frozen at −20 °C until ready for extraction. Remaining 0.5 mL cells were starved for 3 hours in TBST, or metal-chelated TBST, or zinc-supplemented TBST as specified. Following treatment, both pre- and post-starved cells were pelleted and resuspended in 25 DL of 4M formic acid and incubated on ice for 10 minutes. Cell suspensions were frozen on dry ice for 10 minutes and thawed at 37 °C for 10 minutes. The freeze-thaw cycle was repeated 5 times, after which the cell lysates were centrifuged at 13,500 rpm for 5 minutes at 4 °C, and 5 DL of the supernatants containing ^32^P labeled intracellular nucleotides were resolved on a 20 x 20 cm plastic PEI cellulose F TLC plate (Millipore) using 1.5 M KH_2_PO_4_ pH 3.4 as a solvent. When the solvent front reached the top of the plate, the plate was dried and exposed to X-ray film. 5 DL of 100 mM nonradioactive nucleotides [ppGpp (Trilink), ATP, GDP and GTP (Thermo Fisher)] were also resolved as positional markers, which were visualized by handheld UV lamp (Stuart, Cole Palmer). For quantitative analysis, pixel density of ppGpp spot were determined from densitometric scan of radiographs using Fiji (ImageJ). Using the counts from low-zinc culture of wild-type strain in a radiograph as the common denominator, relative counts in other samples from the same radiograph were calculated.

### Cryo-electron microscopy and image processing

Quantifoil holey carbon copper grids R 1.2/1.3 were coated with a continuous layer of carbon (∼50 Å thick). After glow discharge for 30 s on a plasma sterilizer, 4 µl of the 200 nM Rsh-bound pre-elongation complex sample was placed on the grids. The sample was incubated on the grids for 15 s at 4 °C and 100% humidity, followed by blotting for 4 s before flash-freezing into the liquid ethane using a Vitrobot IV (FEI). Data were collected on a Titan Krios electron microscope at 300 keV using a K3 direct electron-detecting camera (Gatan). −1.50 to −2.50 µm defocus range was used at a magnification of 81,000×, yielding a pixel size of 0.846 Å. The dose rate of 23.3 electrons/pixel/second with a 2.05 second exposure time, with 41 frames of 0.05 second duration each, resulted in a total dose of 66.59 e/Å−2. RELION 4.0 was used for data processing. Image stacks were gain-corrected, dose-weighted, and aligned using MotionCor2 (62) for 9,103 micrographs. The contrast transfer function of each aligned micrograph was estimated using CTFFIND-4.1 (63). A subset of micrographs was used for particle picking followed by 2D classification. Relevant 2D classes were used for reference-based particle picking on all micrographs yielding 1,928,609 particles. These particles were 2D classified and suitable class averages selected (389,456 particles) (Fig. S7). The selected particles were 3D refined and followed by 3D classification. Classes corresponding to monosomes with 349,437 particles were selected for further processing. In order to resolve structural heterogeneity in the monosome particles, we performed 3D refinement using a large subunit mask followed by three rounds of fixed orientation 3D classification. In the first round of fixed orientation 3D classification, we used an SSU mask to resolve the SSU conformations into six classes. Class I with 158,489 particles showed density corresponding to P-site tRNA and Rsh, representing the Rsh-bound pre-elongation complex. We then performed the second round of fixed orientation 3D classification on this class using a mask encompassing the entire inter-ribosomal subunit space such as the L7/L12-stalk, A-, P-, and E-site tRNA binding sites, and the L1-stalk regions. The second round of fixed orientation 3D classification resolved the conformational and particularly compositional heterogeneity in the masked region. We derived four classes with P-site tRNA density and two classes with P-site tRNA and Rsh densities. We performed the third round of fixed orientation 3D classification combining the two classes with P-site tRNA and Rsh densities, (49,058 particles) using a mask encompassing the P-site tRNA and Rsh. Two of the classes, one with 16,013 particle and another with 20,308 particles, showed densities corresponding to P-site tRNA and ZBD and ACT domains of Rsh. However, density corresponding to ACT domain was slightly better in the class with 16,013 particles and was refined to a global resolution of 2.98 Å and was used to model the Rsh-ACT domain. Since the densities corresponding to ZBD were very similar in both classes, they were merged. The combined class with a total of 36,321 particles showed significant density corresponding to P-site tRNA and Rsh-ZBD. CTF refinement and Bayesian polishing of the combined class with a total of 36,321 particles yielded a map with a global resolution of 2.7 Å and was used to model the Rsh-ZBD.

### Model building

Coordinates of the large and small subunits from our published C-*M. smegmatis* ribosome structure (PDB:6DZI) were docked as rigid bodies into the cryo-EM map using Chimera 1.14 (64). To achieve optimal fitting, we adjusted the model based on the densities in Coot (65). The model was subsequently refined using PHENIX 1.14 (66). A predicted structure of *M. smegmatis* Rsh is available in the Alphafold Protein Structure database (67). This structure was docked into the corresponding density in the cryo-EM map using Chimera 1.14 (64) and subsequently refined with Coot (65) and PHENIX 1.14 (66). A validation report for the model was obtained from PHENIX 1.14 (66). The overall statistics of EM reconstruction and molecular modeling are listed in Table S3. ChimeraX-1.0 (68) and Chimera 1.14 (64) were used to generate the structural figures in the manuscript.

## Acknowledgments

This work was supported by grants to AKO (NIH: AI132422, NIH: AI163599), CS (AI111696), RKA (NIH:GM61576), and YMH (NIH: GM134931). R.K.A. also acknowledges support to his lab through NIH R01 grants AI132422, GM139277 and AI155473. Support from the Wadsworth Center core facilities–Applied Genomic Technologies and the media core facility– is acknowledged. We also acknowledge Wadsworth Center’s and New York Structural Biology Center’s (NYSBC’s) 3D-EM facilities. NYSBC EM facilities are supported by grants from the Simons Foundation (349247), NYSTAR, the NIH (GM103310) and the Agouron Institute (F00316). Authors acknowledge technical support from Kelley-Hurst, and discussions with Drs. Pallavi Ghosh, Keith Derbyshire, Todd Gray and Erica Lasek-Nesselquist.

## Authors’ Contribution

YL, RT, JC and AKO designed and performed genetics and biochemical experiments; SM, MRS, SRM, NKB and RKA designed and performed structural analysis and model construction; JP, CS, HG and YMH provided reagents. All authors wrote the manuscript.

## Financial Conflict of Interest

Authors declare no financial conflict of interest.

## Data Deposition

The cryo-EM map and atomic coordinates of the *M. smegmatis* 70S-Rsh-fMet-tRNAiMet complex are deposited in the Electron Microscopy and PDB Data Bank (wwPDB.org). Accession codes are EMD-29397 and 8FR8 respectively.

## Supplementary materials

**Figure S1:**
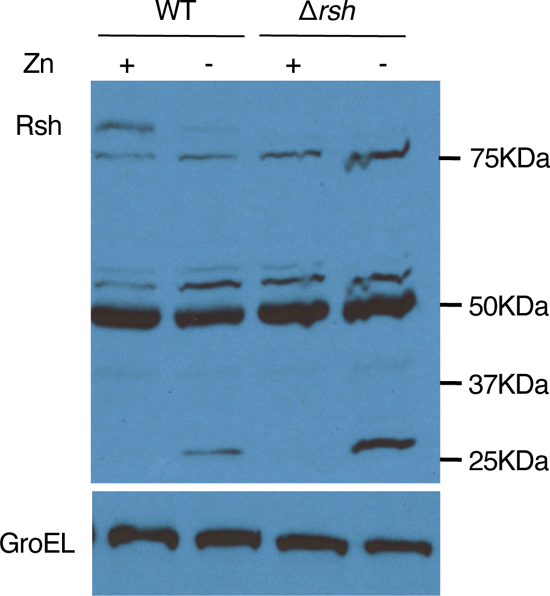
A full-gel immunoblot image of cells lysates from WT and Δ*rsh* strains cultured in high-(1mM ZnSO_4_; abbreviated as Zn) and low-zinc Sauton’s medium (1µM TPEN; abbreviated as T) and probed with anti-Rsh antibody. Data shows lack of any Rsh-derived smaller product in the low-zinc culture. GroEL from the lysate was probed as a loading control.

**Figure S2:**
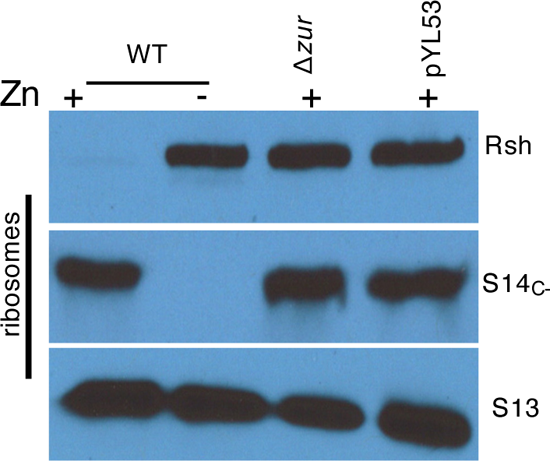
Ribosome remodeling has no effect on Rsh-ribosome interaction. Immunoblot analysis of Rsh bound to 70S ribosomes from 96-hour old high-zinc cultures of WT *M. smegmatis* and its two recombinant strains, Δ*zur* and Δ*c-*:pYL53, which constitutively expressed the remodeled (C-) ribosomes. Low-zinc culture of WT was used as control.

**Figure S3:**
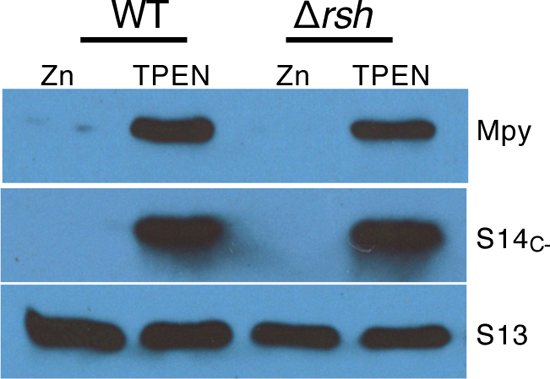
Levels of Mpy in 70S ribosomes purified from high- and low-zinc cultures of WT and Δ*rsh.* S14_c-_ and S13 were probed as controls as indicated. ppGpp

**Figure S4:**
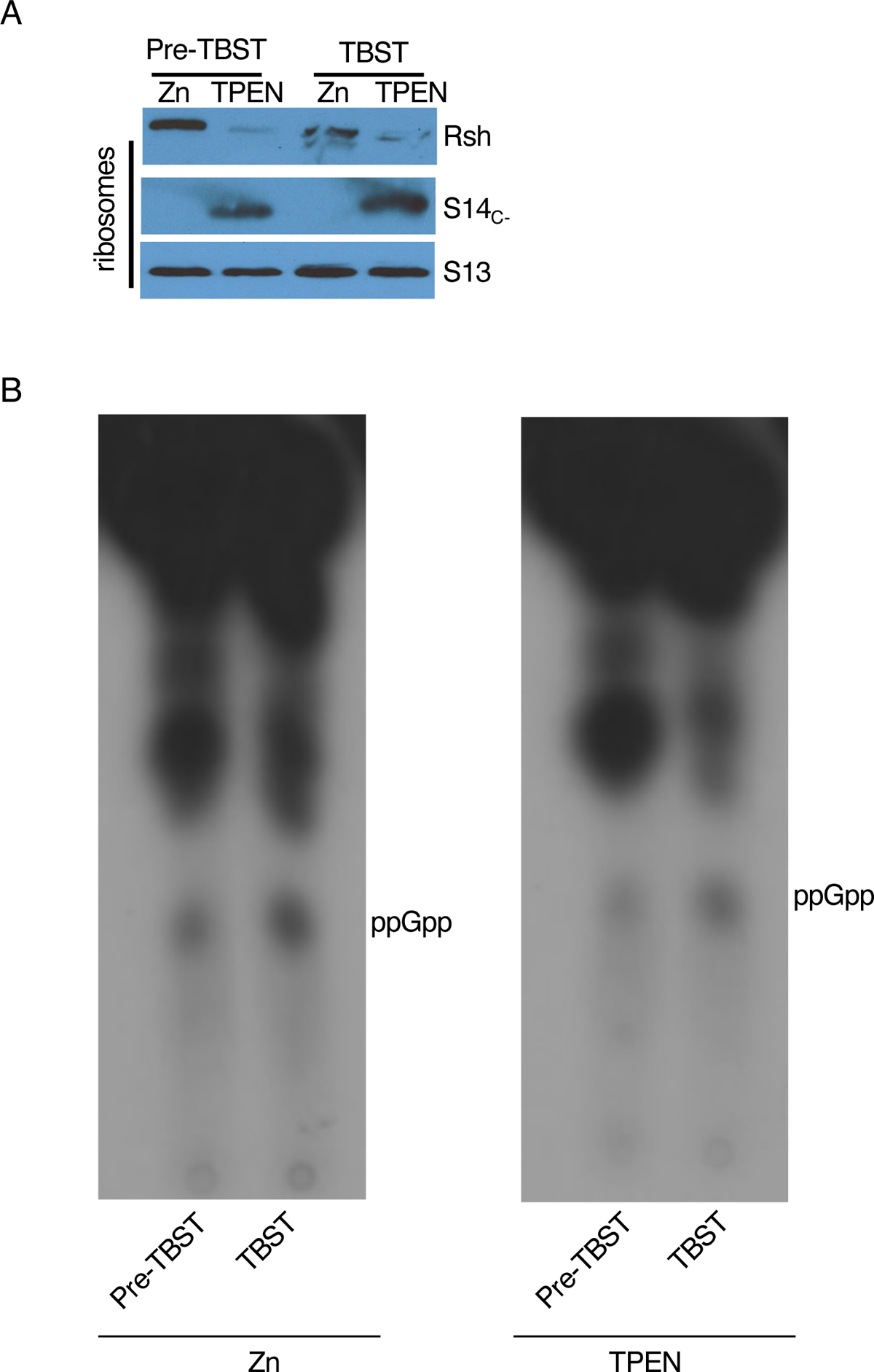
Level of Rsh-bound ribosomes (A) and ppGpp (B) in *M. smegmatis* before and after exposure to starvation (TBST) from high- or low-zinc culture conditions after 28 hours of growth.

**Figure S5:**
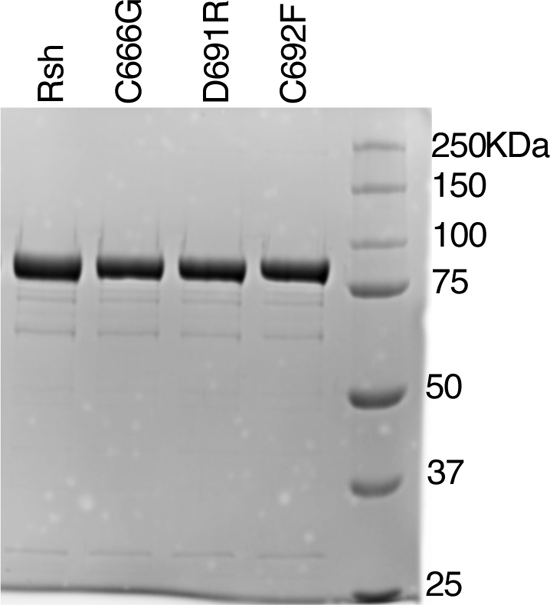
Purified recombinant 6xHis-tagged Rsh proteins expressed from an IPTG-inducible T7 promoter in the *E. coli* BL21 strain and visualized by SDS-PAGE and the Coomassie Blue stain. Each recombinant protein was purified from the same volume of starting cells using the same method, and equal volume of each sample was loaded on the gel.

**Figure S6:**
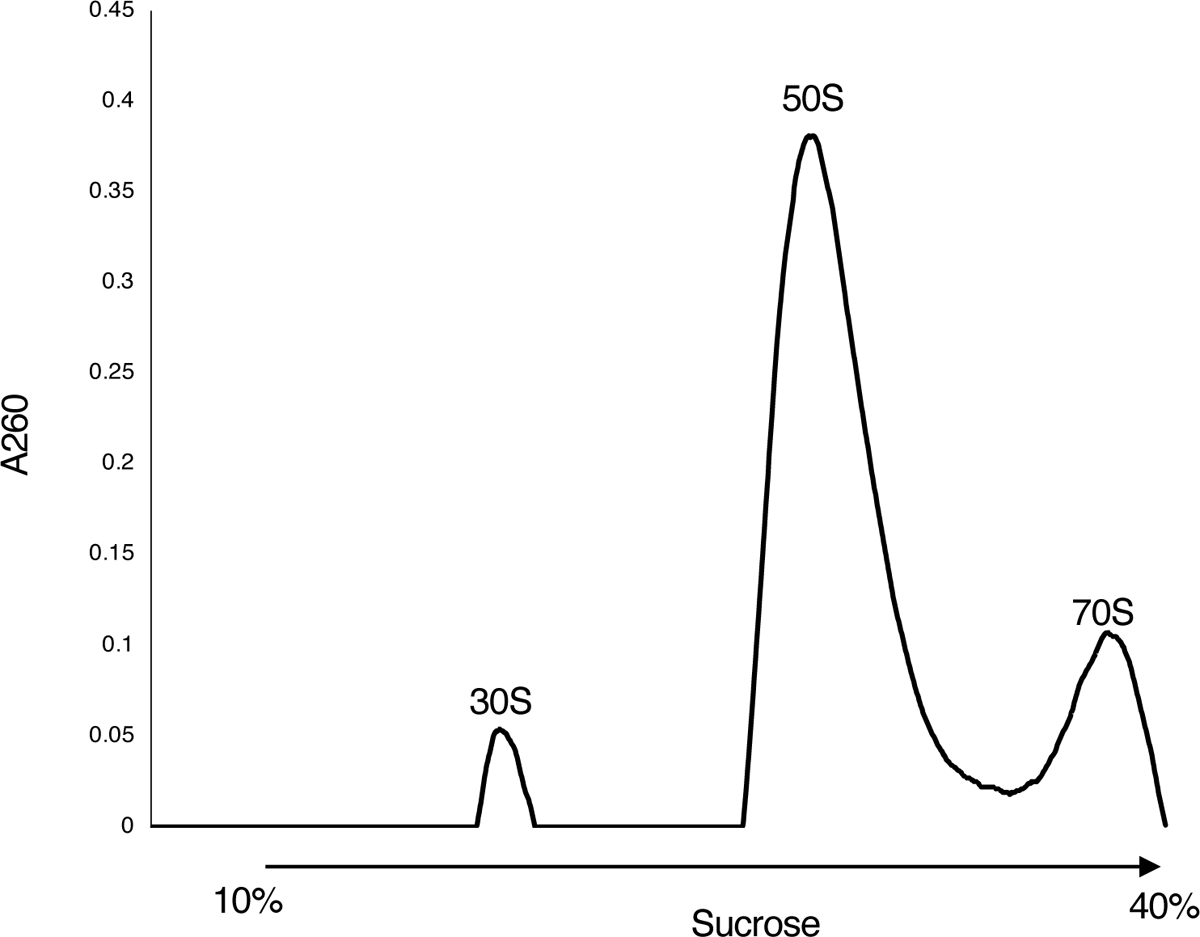
Sucrose density gradient profile of dissociated ribosomes used in the reconstitution experiments described in figures 4B and C. The reconstituted ribosome complex in figure 4C was used for structural studies.

**Figure S7:**
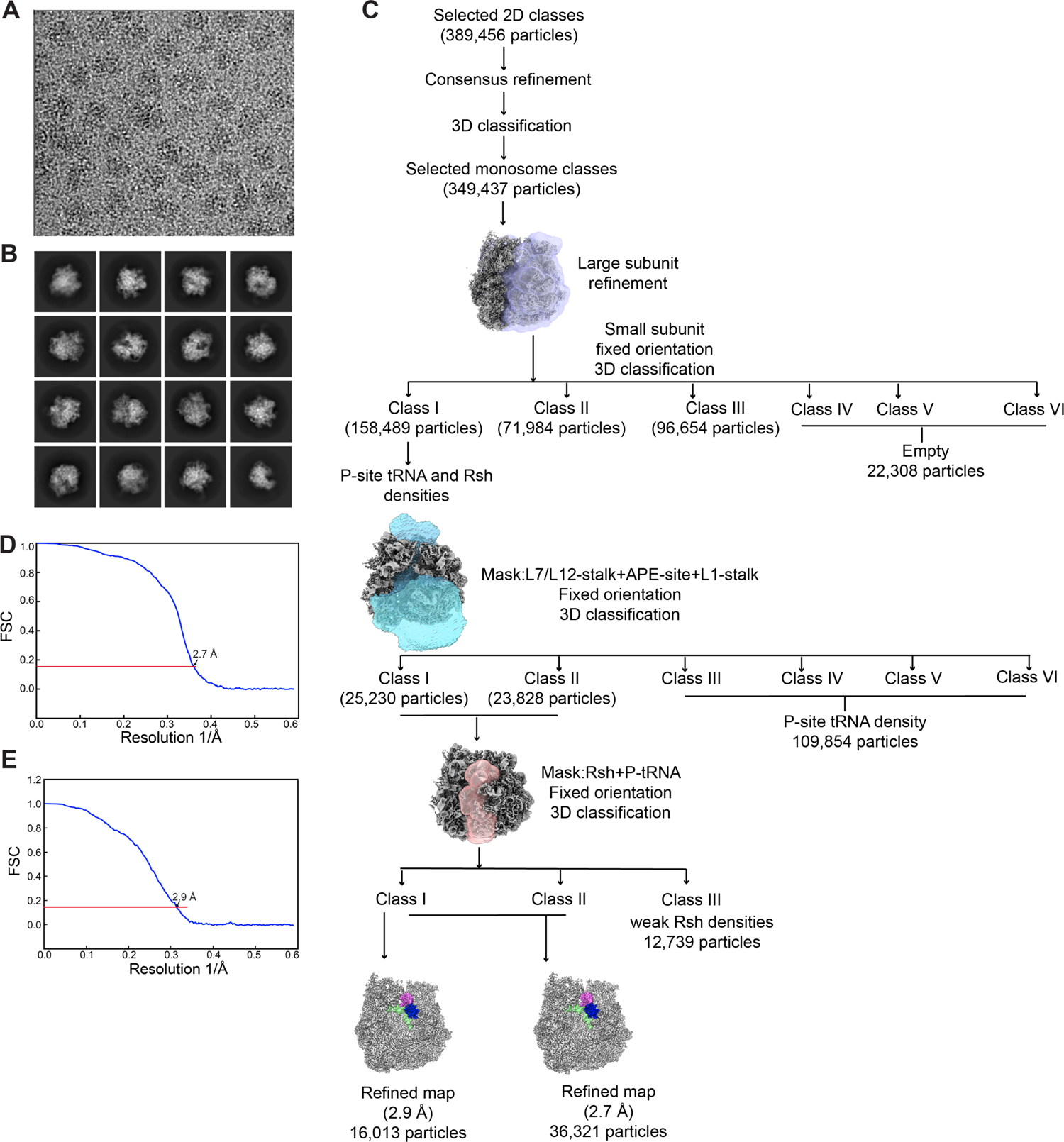
Image processing of M. smegmatis 70S-Rsh-fMet-tRNAiMet complex. **A.** A representative micrograph from the M. smegmatis 70S-Rsh-fMet-tRNAiMet complex dataset. **B.** Representative 2D class averages used for the 3D reconstructions. **C.** Flowchart showing the details of 3D classifications and refinements. **D.** Fourier-shell correlation (FSC) plot for the final map (36,321 particles) used in this study. Local resolution for this map is shown in figure S8. **E.** FSC plot for the map (16,013 particles) used in this study to model the Rsh-ACT domain.

**Figure S8:**
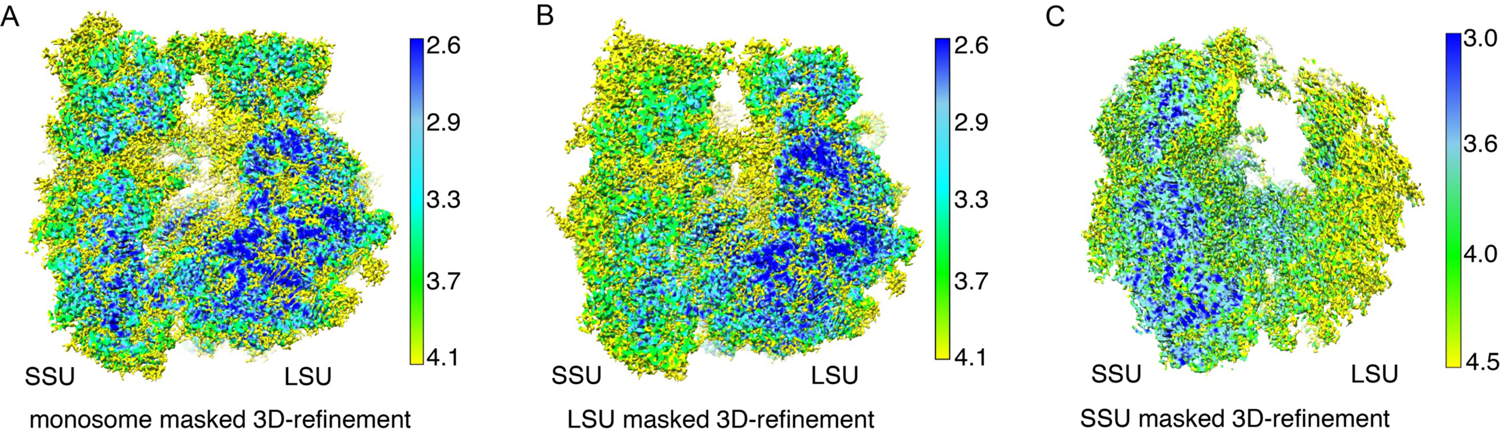
Local resolution of the cryo-EM map of the *M. smegmatis* 70S-Rsh-fMet-tRNAi^Met^ complex. Refined by using (A) 70S monosome, (B) 50S LSU, and (C) 30S SSU masks. In panels B and C, SSU and LSU appear to be partially disordered, as expected. A cutting plane has been applied to show the core of the ribosome in each case.

**Figure S9:**
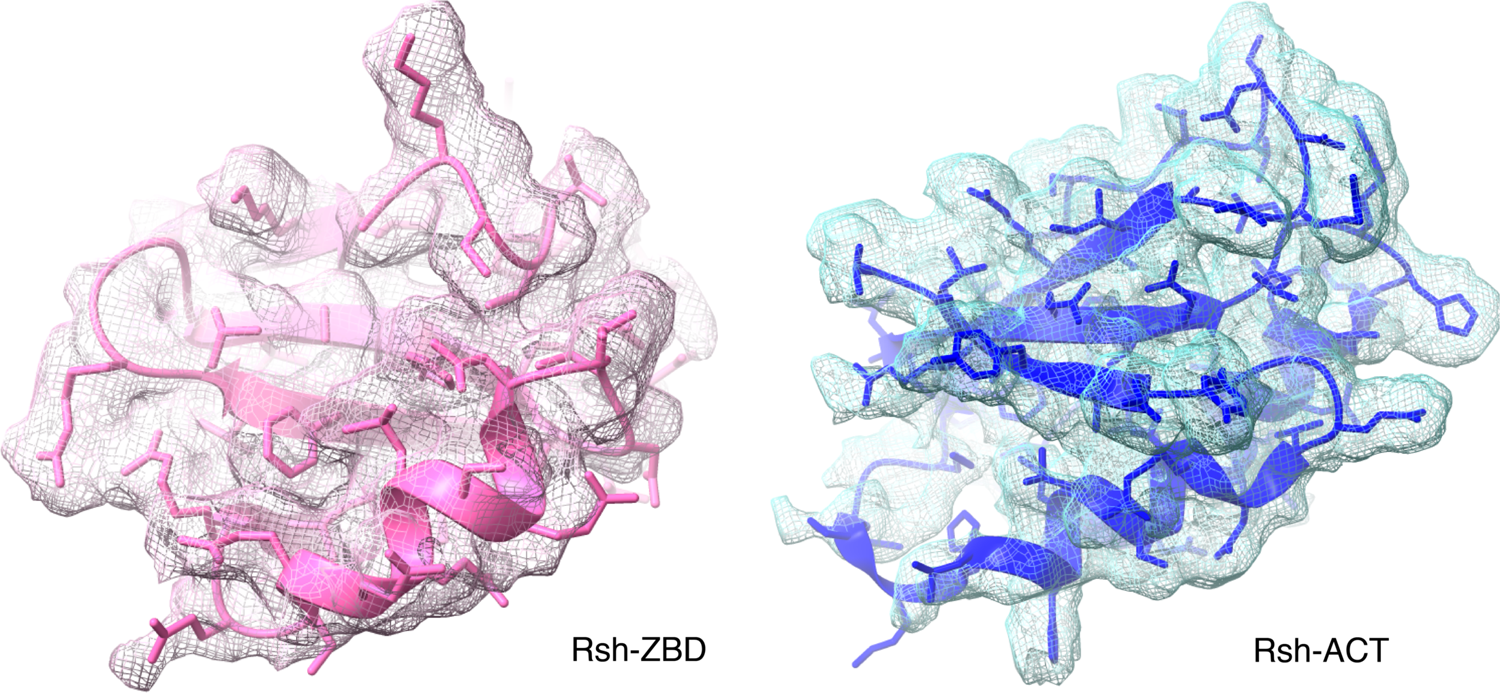
Modeling of the ZB- and ACT domains into corresponding cryo-EM densities. Two different density threshold values were used for modeling the two domains, as density for the main anchoring ZBD (pink) was stronger than that for the ACT domain.

**Figure S10:**
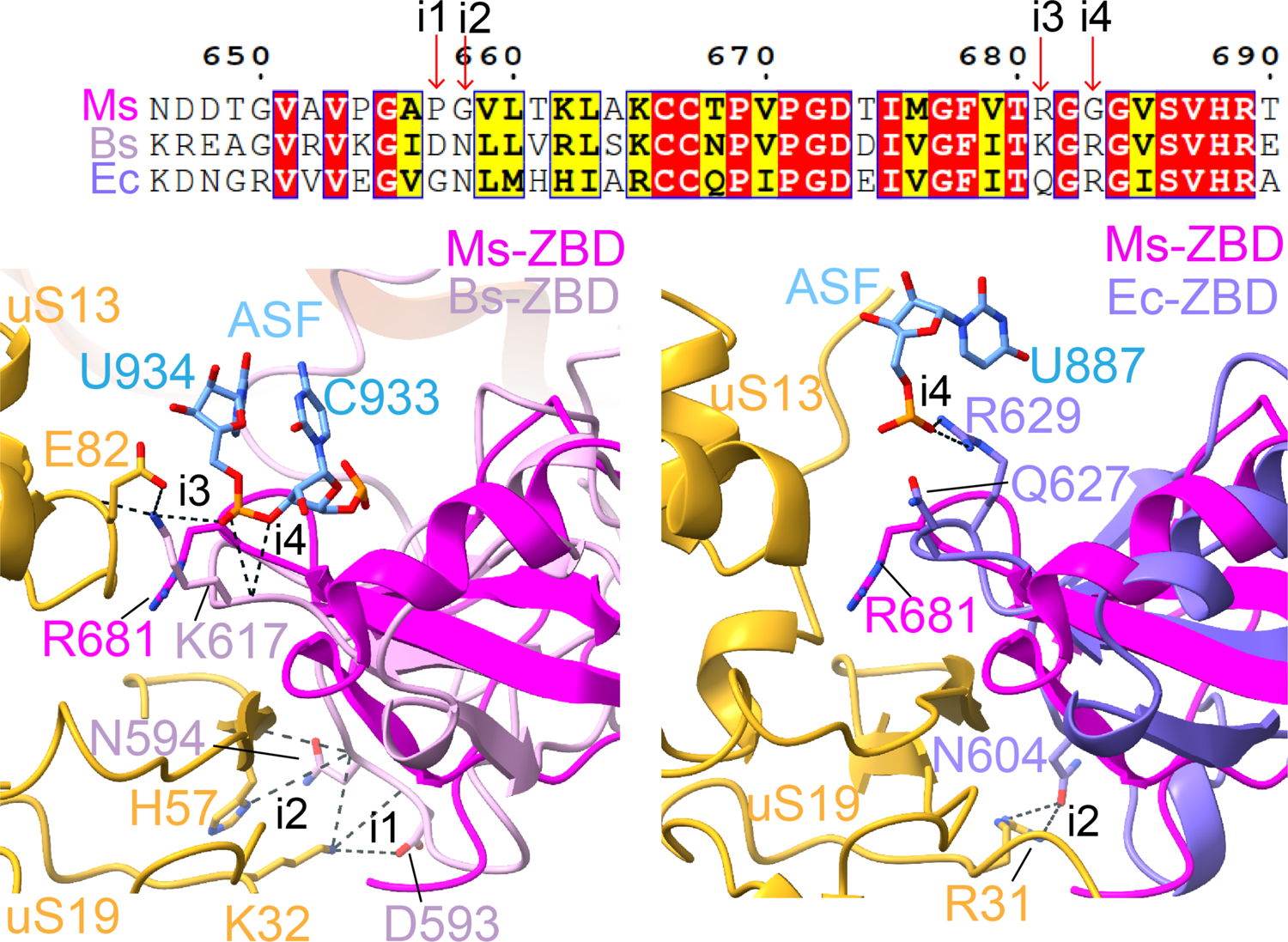
Species-specific differences in Rsh/RelA-ZBD interaction with uS19, uS13, and ASF. Top-Multiple sequence alignment of a short stretch in the ZBD domain of *M. smegmatis* (Ms) Rsh, *B. subtilis* (Bs) Rsh, and *E. coli* (Ec) RelA. Amino acid substitutions at four regions (indicated by red arrows) that result in species-specific differences in interactions (i1-i4) of the ZBD domain with ribosomal components. Bottom left-Superimposition of Ms 70S-Rsh (ZBD, magenta) and Bs 70S-Rsh (ZBD, light pink). Interactions of Bs-Rsh-ZBD with the ribosome that are absent in the case of Ms-Rsh-ZBD are displayed: two with uS19 (i1 and i2), one with uS13 (i3), and one with ASF (i4). Bottom right-Superimposition of Ms 70S-Rsh (ZBD, magenta) and Ec 70S-RelA (ZBD, light purple). Ec RelA-ZBD has two extra interactions with the ribosome, as compared to Ms Rsh-ZBD: One with uS19 (i2) and another with ASF (i4).

---

**Table S1:**
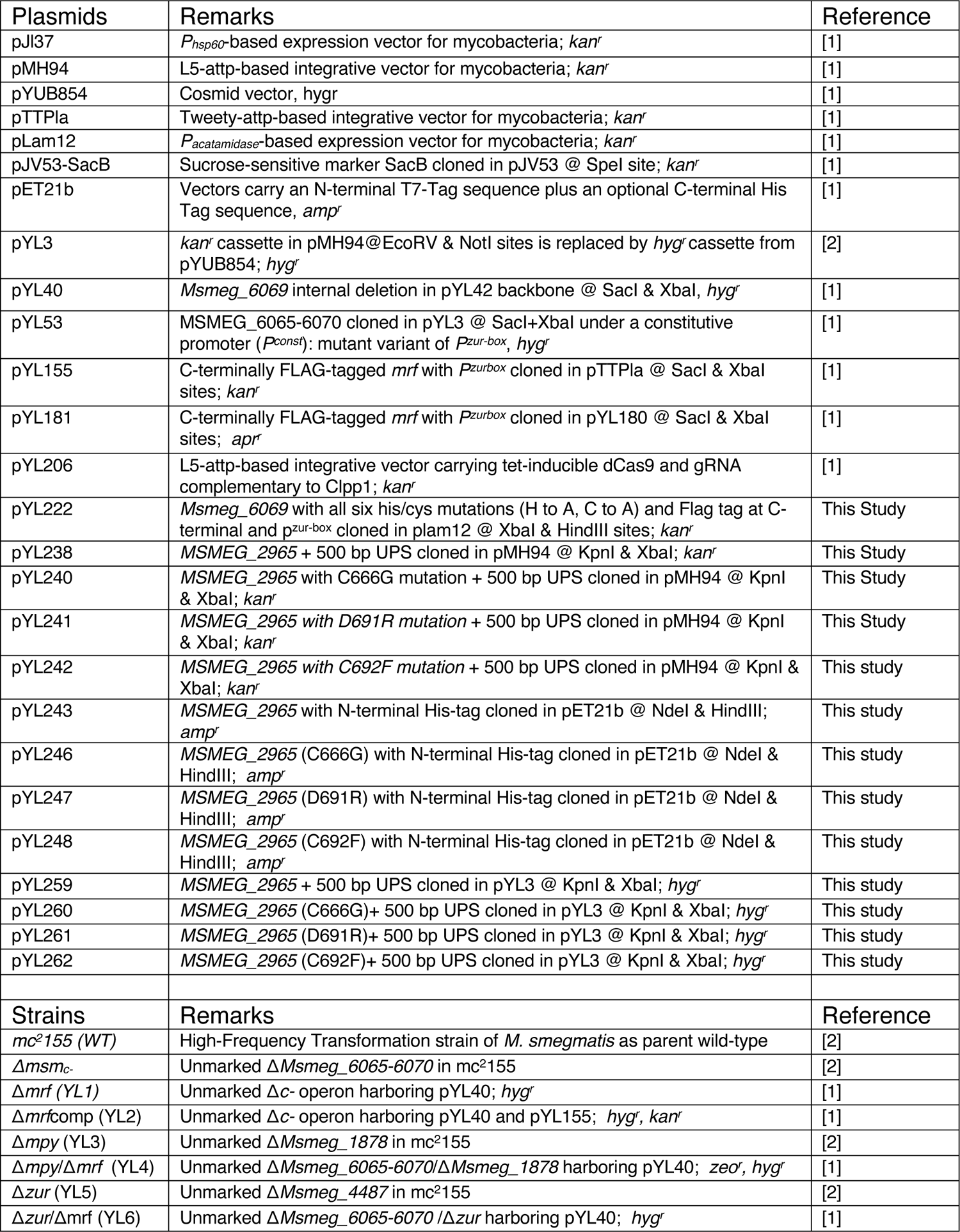
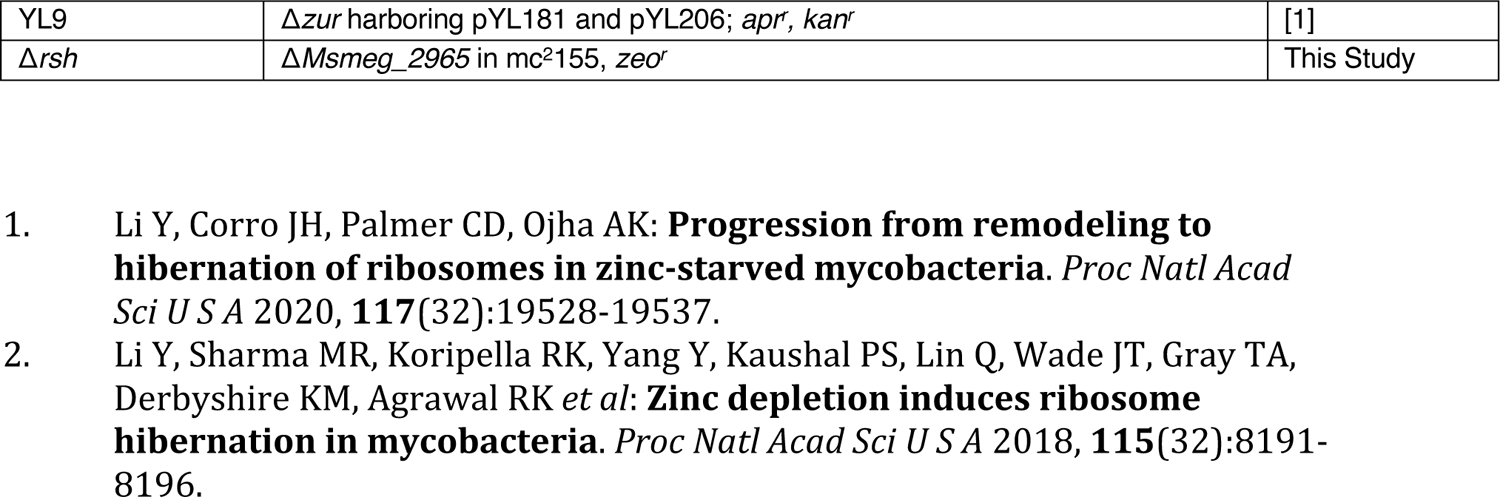
List of plasmids and strains

**Table S2:**
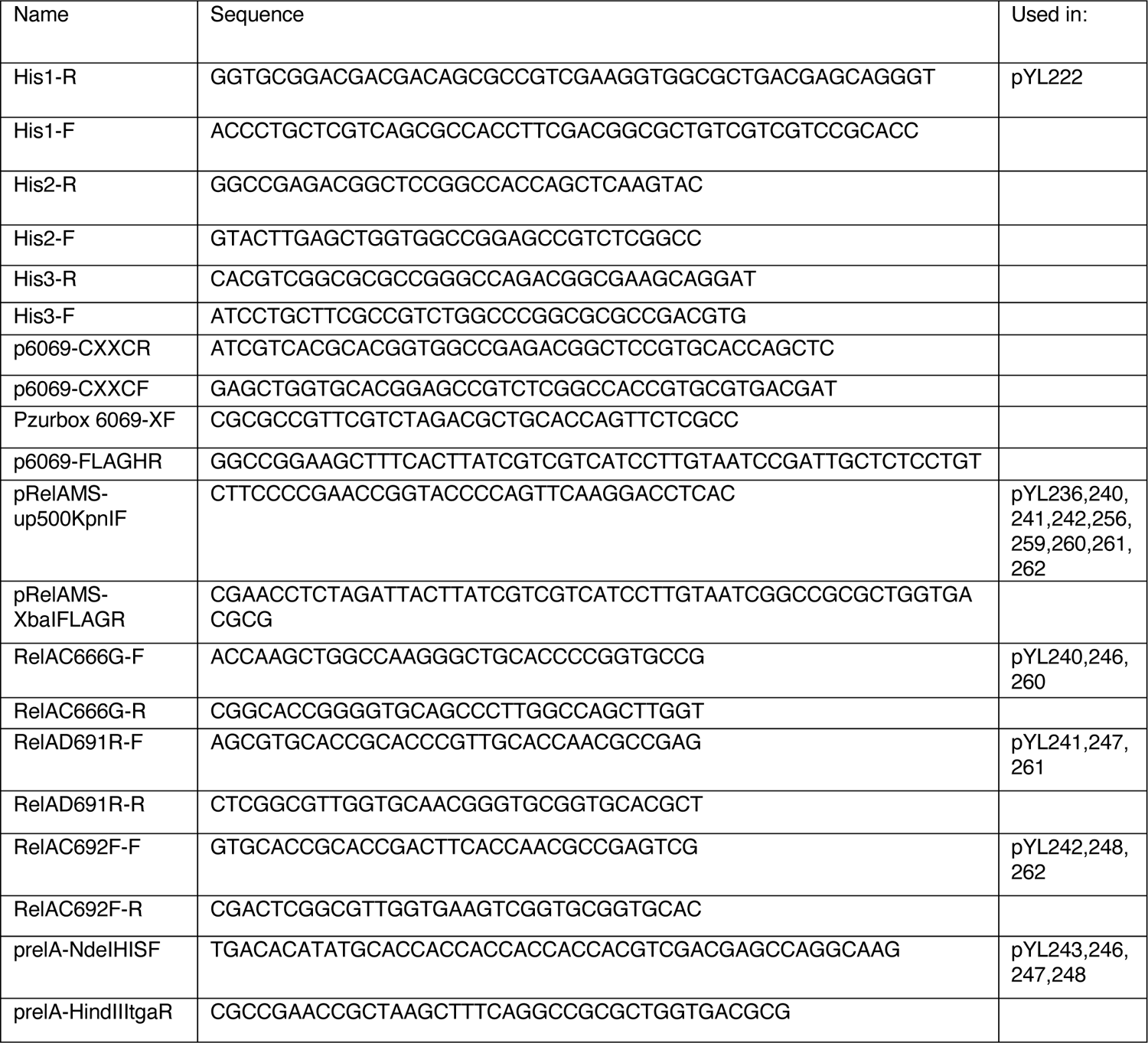
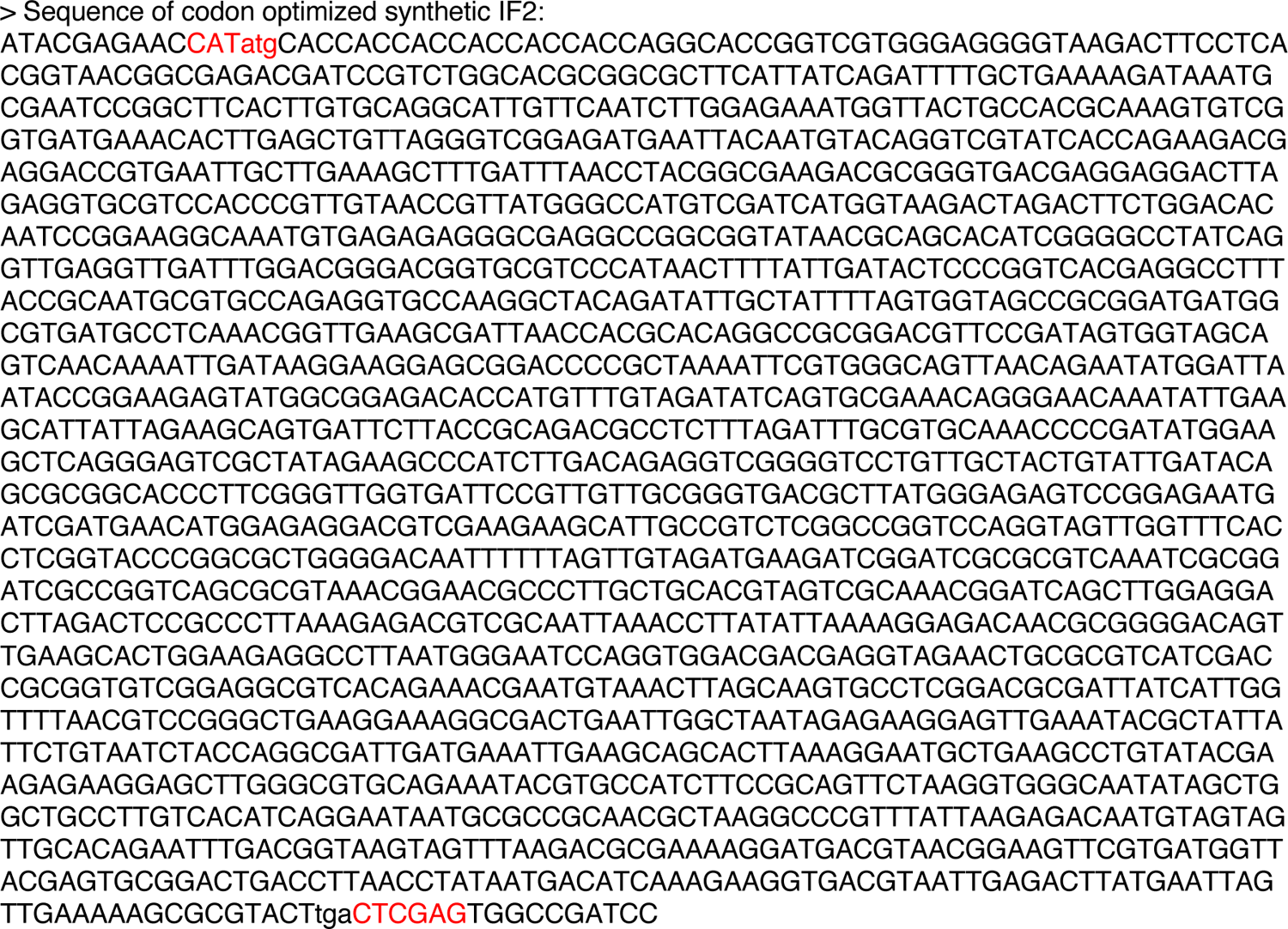
List of oligonucleotides

**Table. S3.** Data collection, Refinement and Model Validation parameters.

| Description | 70S-Rsh-fMet-tRNA<sup>fMet</sup> complex |
| --- | --- |
| **Data collection** |  |
| Microscope | FEI Titan Krios |
| Voltage (kV) | 300 |
| Pixel size (Å) | 0.846 |
| Defocus range (μm) | -1.50 to -2.50 |
| Average e<sup>-</sup> dose per image (e<sup>-</sup>/ Å<sup>2</sup>) | 66.59 |
| Particles (initial) | 1,928,609 |
| Particles (final) | 36,321 |
| FSC-threshold | 0.143 |
| Resolution (Å) | 2.7 |
| Map-sharpening B factor (Å<sup>2</sup>) overall | -44.3 |
| **Refinement** |  |
| RMS deviations |  |
| Bonds (Å) | 0.005 |
| Angles (°) | 0.616 |
| MolProbity score | 2.09 |
| Clash score | 13.51 |
| Rotamer outliers (%) | 0.41 |
| Ramachandran plot |  |
| Outliers (%) | 0.13 |
| Allowed (%) | 6.82 |
| Favored (%) | 93.05 |
| **RNA** |  |
| Correct sugar puckers (%) | 99.24 |
| Angle outliers (%) | 0.00 |
| Bond outliers (%) | 0.00 |
| Good backbone conformations (%) | 78.98 |
| Model composition |  |
| RNA bases | 4843 |
| Protein residues | 6299 |

## Notes

### Competing Interest Statement

The authors have declared no competing interest.

### Summary of Updates

This version of the manuscript includes new analyses and interpretations of the data.

